# Protein-protein interactions in the Mla lipid transport system probed by computational structure prediction and deep mutational scanning

**DOI:** 10.1101/2022.12.09.519820

**Authors:** Mark R. MacRae, Dhenesh Puvanendran, Max A. B. Haase, Nicolas Coudray, Ljuvica Kolich, Cherry Lam, Minkyung Baek, Gira Bhabha, Damian C. Ekiert

**Affiliations:** Department of Cell Biology, New York University School of Medicine, New York, United States; Applied Bioinformatics Laboratories, New York University School of Medicine, New York, United States; Department of Biochemistry, University of Washington, Seattle, WA, USA; Institute for Protein Design, University of Washington, Seattle, WA, USA; Department of Microbiology, New York University School of Medicine, New York, United States

## Abstract

The outer membrane (OM) of Gram-negative bacteria is an asymmetric bilayer that protects the cell from various external stressors, such as antibiotics. The Mla transport system is implicated in the <u>M</u>aintenance of outer membrane <u>L</u>ipid <u>A</u>symmetry and is thought to mediate retrograde phospholipid transport across the cell envelope. This system uses a shuttle-like mechanism to move lipids between the MlaFEDB inner membrane complex and the MlaA-OmpF/C OM complex, via a periplasmic lipid-binding protein, MlaC. MlaC binds to MlaD and MlaA, but the underlying protein-protein interactions that facilitate lipid transfer are not well understood. Here, we take an unbiased deep mutational scanning approach to map the fitness landscape of MlaC, which provides insights into important functional sites. Combining this analysis with AlphaFold2 structure predictions and binding experiments, we map the MlaC-MlaA and MlaC-MlaD protein-protein interfaces. Our results suggest that the MlaD and MlaA binding surfaces on MlaC overlap to a large extent, leading to a model in which MlaC can only bind one of these proteins at a time. Low-resolution cryo-electron microscopy (cryo-EM) maps of MlaC bound to MlaFEDB suggest that at least two MlaC molecules can bind to MlaD at once, in a conformation consistent with AlphaFold2 predictions. These data lead us to a model for MlaC interaction with its binding partners and insights into lipid transfer steps that underlie phospholipid transport between the inner and outer membranes.

## Introduction

The outer membrane (OM) of Gram-negative bacteria acts as a protective barrier against various harmful environmental factors, including antibiotics. The OM is an asymmetric bilayer, composed of phospholipids (PLs) in the inner leaflet and lipopolysaccharide (LPS) in the outer leaflet, and is separated from the inner membrane (IM) by an aqueous periplasmic space. The asymmetry of the OM is important for creating a membrane that is largely impermeable to both hydrophilic and hydrophobic molecules. To maintain the asymmetry of the OM and the integrity of the OM barrier, phospholipids must be moved between the OM and IM, across the periplasm. The <u>M</u>aintenance of OM <u>L</u>ipid <u>A</u>symmetry (Mla) system is perhaps the best-understood phospholipid transporter in Gram-negative bacteria, and is thought to have a key role in maintaining the OM by removing mislocalized PLs from the outer leaflet and transporting them to the IM for recycling (Abellón-Ruiz et al., 2017; Chong et al., 2015; Ekiert et al., 2017; Ercan et al., 2019; Hughes et al., 2019; Isom et al., 2017; Kamischke et al., 2019; Malinverni and Silhavy, 2009; Powers and Trent, 2018; Shrivastava and Chng, 2019; Thong et al., 2016; Yeow et al., 2018).

The Mla system consists of three main parts: 1) MlaFEDB in the IM, an ABC (ATP Binding Cassette) transporter complex (Chi et al., 2020; Coudray et al., 2020; Ekiert et al., 2017; Kamischke et al., 2019; Mann et al., 2021; Tang et al., 2021; Thong et al., 2016; Zhang et al., 2020; Zhou et al., 2021), 2) MlaC, a periplasmic lipid trafficking protein (Ekiert et al., 2017; Ercan et al., 2019; Hughes et al., 2019), and 3) MlaA-OmpF/C, an OM complex (Abellón-Ruiz et al., 2017; Chong et al., 2015; Yeow et al., 2018). To maintain the asymmetry of the OM, the current model for Mla function suggests that it most likely carries our retrograde PL transport (Low et al., 2021; Tang et al., 2021): First, mislocalized PLs in the outer leaflet of the OM are recognized by MlaA-OmpF/C and can cross the OM via a channel through the center of MlaA (Abellón-Ruiz et al., 2017). MlaC binding to MlaA facilitates transfer of PLs to MlaC, which acts as a ferry to transport PLs across the periplasm. The lipid cargo is bound in a hydrophobic cavity between the β-sheet and α-helical subdomains of MlaC (**Fig. 1a**) (Ekiert et al., 2017), via a proposed clamshell mechanism (Hughes et al., 2019). After traversing the periplasm, MlaC binds to the MlaFEDB complex in the IM (Ekiert et al., 2017; Hughes et al., 2019), which accepts the PL from MlaC and uses ATP hydrolysis to translocate the PL into the IM. Structures of MlaC in the apo (Hughes et al., 2019) and lipid-bound (Ekiert et al., 2017) states, together with biochemical studies (Ercan et al., 2019; Yeow and Chng, 2022) have begun to shed light on MlaC function.

**Figure 1.**
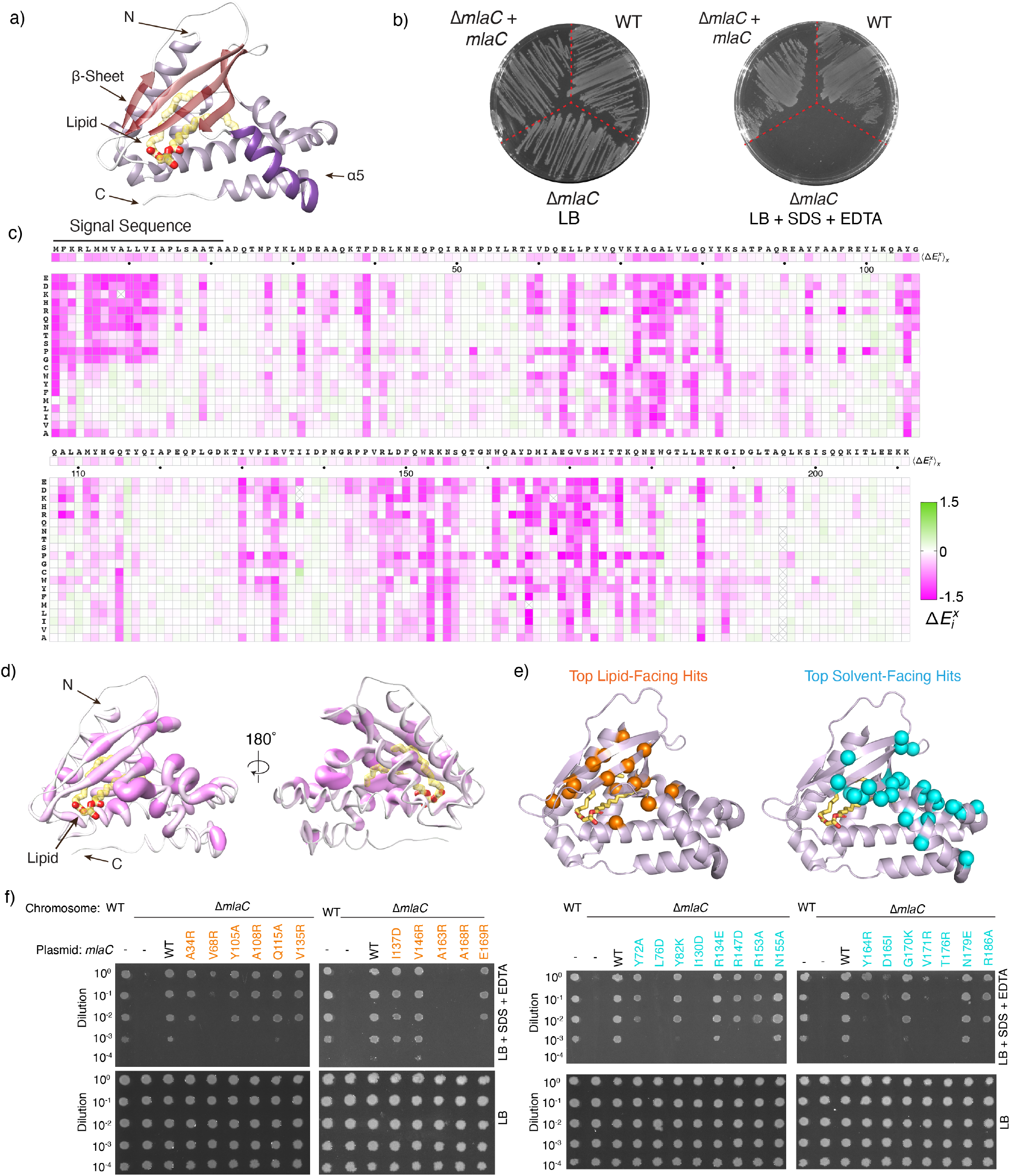
Deep mutational scanning and Identification of functionally important regions of MlaC. **a** Crystal structure of MlaC (PDB 5UWA) in cartoon representation with β-sheet in red and ɑ5 helix in purple. A phospholipid bound in the pocket of MlaC is shown as yellow sticks. **b)** Plating of *E. coli* strain BW25113 (WT), the *mlaC* deletion mutant (Δ*mlaC*), and Δ*mlaC* complemented with WT *mlaC* on a plasmid (Δ*mlaC* + *mlaC*) on LB agar with or without SDS+EDTA. Both plates also include 2% arabinose to induce mlaC expression from the plasmid. **c)** Heat map summarizing results of deep mutational scanning of MlaC (average of 2 biological replicates). X-axis: the sequence of WT MlaC from N-terminus to C-terminus; Y-axis: all possible amino acid substitutions. Each square represents the fitness cost of an individual mutation (Δ*E*_*i*_^*x*^) relative to WT. Mutations that decrease fitness relative to the WT are shown in shades of magenta, while mutations that increase fitness are in shades of green, with white representing neutral mutations, as shown in the key. The mean effect of mutations on fitness at each position is shown in the horizontal strip just below the MlaC sequence. Boxes marked with “X” indicate no coverage in the library. **d)** MlaC structure oriented roughly as in (a) with residues colored by mean fitness cost of mutations as in (c); radius of each residue corresponds to the fitness key, with larger radius corresponding to the most decreased fitness. Residues with most decreased fitness appear as the deepest shade of magenta and with the largest radius. Most functionally important residues map towards one face of MlaC (left), while the opposite face is less sensitive to mutation (right). **e)** Many of the residues with the most decreased fitness from deep mutational scanning of MlaC map to the lipid-binding pocket (left; orange spheres; 15 residues) or a solvent-accessible cluster (right; cyan spheres; 25). **f)** Genetic complementation of Δ*mlaC E. coli* cells with plasmids expressing WT MlaC or point mutations in the lipid binding pocket (left, orange) or on the solvent-accessible surface (right, cyan). 10-fold serial dilutions of strains were spotted on LB agar with or without SDS+EDTA. All plates contain 2% arabinose to induce *mlaC* expression from plasmid.

Based on the current model of lipid transport by the Mla system, MlaC must bind to MlaA and MlaD in order to transfer lipids, but a structural understanding of these interactions remains elusive. Challenges to obtaining high-resolution structures of MlaC-MlaD and MlaC-MlaA complexes include the low binding affinity, as well as compositional and conformational heterogeneity. In this study, we use deep mutational scanning to map the functionally important regions of MlaC and subsequently use a combination of biochemical assays, structural approaches, and computational structure prediction to suggest how MlaC interacts with both MlaD and MlaA. Our results provide insights into protein-protein interactions and lipid transfer steps within the Mla transport pathway in *E. coli*.

## Results

### Deep mutational scanning identifies functionally important residues in MlaC

To gain unbiased insight into functionally important residues in MlaC, we performed deep mutational scanning (Fowler and Fields, 2014). We constructed a pooled plasmid library of 4,431 unique *mlaC* variants, where the codon for each residue was mutated to every other possible amino acid (20 aa + stop at 211 positions). This library was transformed into an *E. coli mlaC* knockout strain, which is unable to grow on LB agar supplemented with SDS and EDTA, but can be complemented by WT MlaC on a plasmid (Ekiert et al., 2017; Malinverni and Silhavy, 2009). Using Next-Generation Sequencing as a readout, we measured the relative frequency of each *mlaC* mutant in the pool after selection for growth in the presence of SDS+EDTA vs. non-selective medium (**Fig. 1b**). We assessed the impact of each mutation, resulting in a comprehensive fitness landscape for MlaC (**Fig. 1c**). For each mutation, we calculated a relative fitness value (Δ*E*_*i*_^*x*^) indicating the mutational tolerance at each residue of MlaC. Functionally important residues in MlaC are less tolerant of mutations and therefore are expected to have negative fitness values. Nearly all mutations in MlaC were either neutral or negatively impacted protein fitness; we did not identify any mutations in MlaC that significantly increased SDS+EDTA resistance above the level of the parent, WT *E. coli* strain.

We selected for further analysis MlaC residues (Supplementary Data File 1) for which 5 or more mutations resulted in a fitness change of more than one standard deviation from the mean fitness of all mutations. The resulting 59 residues can be divided into four functionally distinct groups: 1) residues in the signal peptide, which directs the secretion of MlaC to the periplasm (12 residues, including initiator Met); 2) buried residues that may be important for protein stability (7 residues); 3) residues lining the lipid-binding pocket, likely important for lipid interaction (15 residues); and 4) residues on the solvent-exposed surface without obvious functional significance (25 residues). While the importance of the first three groups of residues can be readily understood, the role of the final group of surface-exposed residues is less clear. We hypothesized that these may be involved in protein-protein interactions with MlaA or MlaD, or other functions unrelated to lipid binding. Most of these surface-exposed residues are clustered on one side of MlaC, suggesting that this surface is functionally important (**Fig. 1d and 1e**). Based on a combination of mutational fitness cost and location in the MlaC structure, we selected a panel of residues located within the lipid binding pocket and solvent exposed regions to validate our deep mutational scanning (**Fig. 1e**). To assess the function of MlaC mutants, we used a cell growth assay based upon genetic complementation of *E. coli mla* knockout strains. Strains with single deletions of *mlaC, mlaA*, or *mlaD* are unable to grow in the presence of SDS and EDTA, but growth can be rescued by a functional copy of the missing gene on a plasmid (**Fig. 1f**). Thus, we constructed a series of plasmids expressing MlaC mutants, and assessed their ability to restore the growth of an *mlaC* knockout strain in the presence of SDS and EDTA. From the lipid-facing subgroups, the A163R and A168R mutations in MlaC had the strongest effect, failing to complement an *mlaC* knockout strain. From the solvent-facing subgroup, L76D, I130D, D165I, V171R and T176R had the strongest effect (**Fig. 1f**). Other mutations have milder or undetectable effects on cell growth. All of the selected lipid and solvent facing mutants are expressed at similar levels in cell lysates, suggesting that differences in apparent fitness are not due to altered protein expression (**Fig. S1e**). The complementation experiments are generally in agreement with the results of deep mutational scanning. However, some mutations with significant fitness defects from deep mutational scanning show little impact on growth in the complementation assay. This may reflect greater sensitivity of deep mutational scanning to identify mutations moderately impacting MlaC function. Together, these data suggest that clustered residues on the surface of MlaC are important for its function, and could potentially represent a protein-protein interaction interface.

### AlphaFold2 prediction of MlaC-MlaA complex is consistent with mutational analysis

MlaC has been shown to interact with MlaA and MlaD *in vitro* (Ekiert et al., 2017; Ercan et al., 2019). We hypothesized that some of the surface-exposed residues identified by deep mutational scanning of MlaC may be important for these interactions. AlphaFold2 (Jumper et al., 2021) has emerged as a powerful tool for the prediction of protein-protein complexes, especially in prokaryotes, and can be used in combination with experiments to validate and probe structural interactions. We predicted the structure of the MlaC-MlaA complex using AlphaFold-Multimer (Evans et al., 2022) as implemented on COSMIC^2^ (Cianfrocco et al., 2017; Cianfrocco MA, Wong M, Youn C, Wagner R, Leschziner AE, n.d.)(**Fig. 2a**). All five of the highest ranked predictions for the MlaC-MlaA complex are nearly identical to one another (**Fig. S2a and S2b**), which may indicate a reliable prediction with a rigid interaction interface. As expected, the individual MlaC and MlaA components of the complex are consistent with previously published structures (Abellón-Ruiz et al., 2017; Ekiert et al., 2017) (**Fig. S2c and S2d**). However, the C-terminal region of MlaA (residues 226-251), which was not resolved in previous X-ray structures (Abellón-Ruiz et al., 2017), is predicted to extend away from the membrane into the periplasm where it interacts with MlaC (**Fig. 2a**). Our predictions appear to be globally similar to other recently reported predictions (Yeow and Chng, 2022).

**Figure 2.**
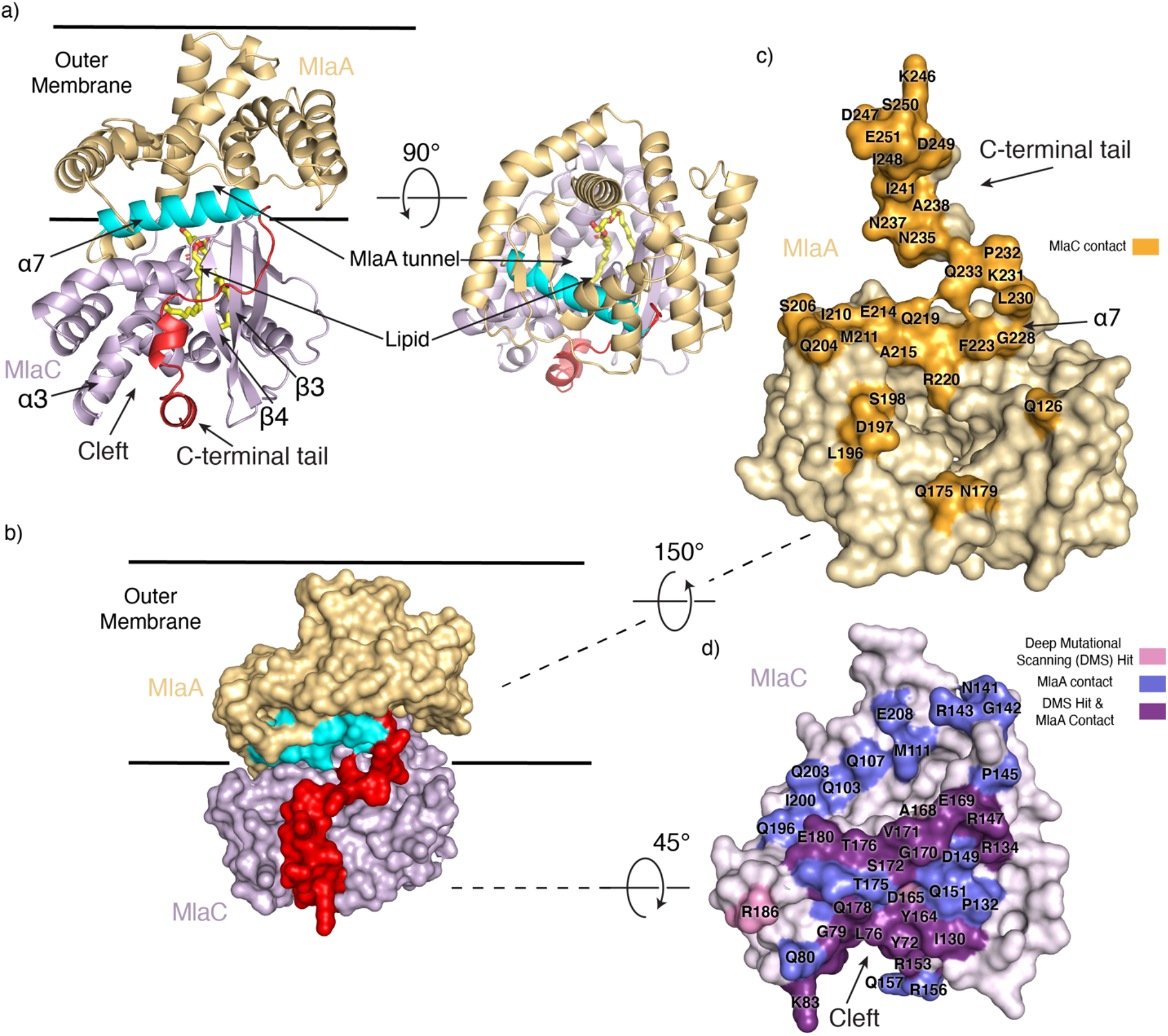
AlphaFold Multimer prediction of MlaC-MlaA protein-protein interface. **a)** AlphaFold Multimer prediction of MlaC (purple) bound to MlaA (gold). The C-terminal tail of MlaA (red) wraps around MlaC and binds in a cleft formed by helix ɑ3 and strands β3 and β4 on MlaC.The ɑ7 helix of MlaA (cyan) is also a key part of the predicted interface. This predicted binding mode aligns the channel of MlaA with the lipid-binding pocket of MlaC. The lipid was not included in AlphaFold multimer predictions, but is shown based on its location in the crystal structure (5UWA). **b)** The predicted MlaC-MlaA model shown in surface representation. **c)** Surface representation of MlaA; residues in MlaA predicted to be in contact with MlaC are colored in bright orange and labeled. **d)** Surface representation of MlaC; Residues in MlaC predicted to be within 4 Å distance of MlaA from AlphaFold Multimer, shown to have reduced fitness from deep mutational scanning (DMS), or both, are mapped onto the structure as indicated in the key.

The AlphaFold-multimer model suggests that 44 MlaC residues and 37 MlaA residues may be within 4 Å of each other and could be involved in interactions within the complex (**Fig. 2b, 2c and 2d**). The prediction shows that residues at the rim of the MlaA channel may interact with residues surrounding the lipid binding pocket on MlaC (**Fig. 2a, 2c and 2d**). This mode of interaction results in the lipid being poised for transfer between MlaA and MlaC. Moreover, among the 25 solvent-exposed residues of MlaC chosen for further analysis from deep mutational scanning, 21 of them lie at the predicted interface with MlaA and can be divided into 2 distinct groups (**Supplementary Data 1 and Fig. 2d**). First, 15 of these MlaC residues are positioned to interact with the C-terminal tail of MlaA, which wraps around the side of MlaC and docks two short helices in a cleft on MlaC formed between helix ɑ3 and strands β3/β4. Second, the remaining 6 MlaC residues are predicted to interact with the ɑ6 helix on MlaA.

To assess the functional importance of MlaA residues predicted to interact with MlaC, we designed a panel of mutations in MlaA. Using the cell growth assay, we tested whether these mutants could genetically complement an *E. coli mlaA* knockout strain, which is unable to grow in the presence of SDS and EDTA unless a functional copy of *mlaA* is provided on a plasmid (**Fig. 3c**). First, we designed three mutants in which the C-terminal region of MlaA was truncated to varying degrees: MlaA(Δ244-251), MlaA(Δ238-251) and MlaA(Δ227-251) (**Fig. 3a**). All three mutants are unable to support growth under these conditions, similar to the *mlaA* knockout. Second, we generated point mutations in the C-terminal helices, I241N, L245N and I248N (**Fig. 3b**); all three point mutations do not grow in our complementation assay. Similar truncations of the C-terminal tail of MlaA were recently reported (Yeow and Chng, 2022), as well as point mutations in charged residues, which also showed decreased growth in the presence of SDS and EDTA. Third, we designed three single mutants on the rim of the tunnel through MlaA, F223N, L230 and D198N (**Fig. 3b**): F223N does not grow in our complementation assay, while L230N and D198N grow similarly to WT MlaA (**Fig. 3c**).

**Figure 3.**
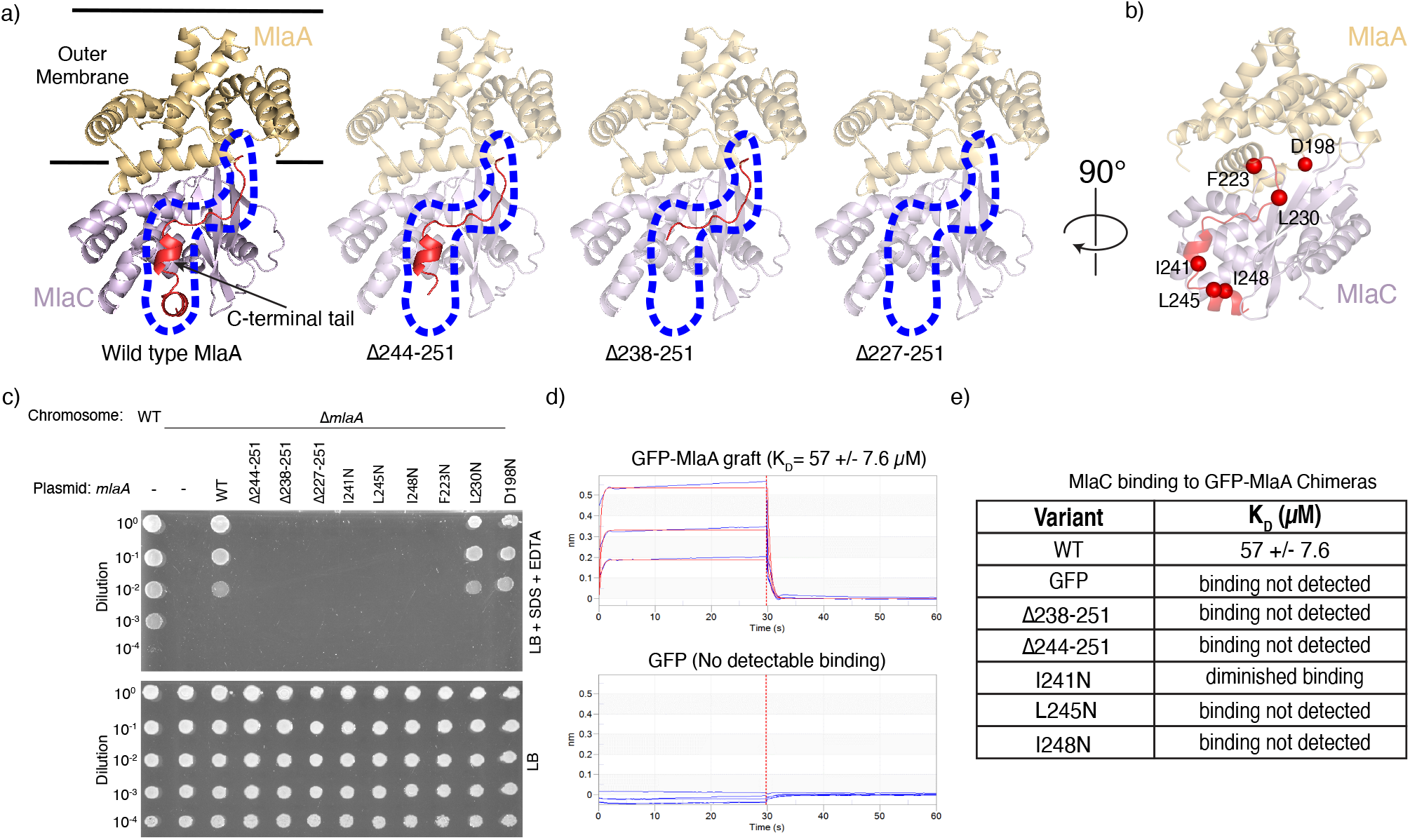
Functional assays probing the role of the MlaA C-terminal tail for interaction with MlaC. **a)** Truncations of varying lengths of the MlaA C-terminal tail (red) used for genetic complementation assays. The area corresponding to the C-terminal tail in the WT prediction is outlined with a blue dotted line. **b)** Residues in MlaA selected for genetic complementation assays. **c)** Genetic complementation of Δ*mlaA E. coli* cells with plasmids expressing WT MlaA, point mutations, or C-terminal truncations. 10-fold serial dilutions of strains were spotted on LB agar with or without SDS+EDTA as indicated. All plates contain 2% arabinose to induce *mlaA* expression from plasmid. **d)** Binding curves from biolayer interferometry show association and dissociation of MlaC to either the GFP-MlaA C-terminal tail chimeric mutant (apparent K_D_ = 57 +/-7.6 μM) or GFP only, as a negative control (no detectable binding). Binding curves show association signal (nm) on the Y-axis and time (s) on the X-axis. A 1:1 binding model is shown (red) for varying concentrations of MlaC (blue). **e)** Summary from binding experiments of WT MlaC to different GFP-MlaA C-terminal tail chimera variants.

Since many mutations affecting the function of MlaC and MlaA map to the predicted interaction interface involving the MlaA C-terminal tail, we hypothesized that the MlaA C-terminal tail may be a major binding determinant for MlaC. To test this hypothesis, we grafted the C-terminal tail of MlaA (residues 227-251) onto a neutral scaffold protein (GFP), and assessed whether MlaC was capable of binding to this chimeric protein [GFP-MlaA(227-251)] using biolayer interferometry. MlaC binds to the GFP-MlaA(227-251) chimera with a dissociation constant (*K*_*D*_) of 57 +/-7.6 μM, while binding of MlaC to GFP alone (negative control) was not detected (**Fig. 3d**). Thus, MlaC is capable of binding to the C-terminal tail of MlaA in isolation, though it is unclear how the affinity of this interaction compares to the affinity of MlaC for native MlaA, which is currently unknown. By making a series of mutations within this chimeric GFP-MlaA(227-251) framework, we were able to test how the point mutations and truncations tested in the cell growth assay above impact MlaC binding. In good agreement with the results of our complementation experiments, we found that MlaC is unable to bind to the Δ244-251 and Δ238-251 truncations; the point mutations L245N and I248N also abolished binding. For the final point mutant, I241N, binding to MlaC is diminished (**Fig. 3e and S3**). These results suggest that the C-terminal tail of MlaA plays an important role in MlaC binding, as point mutations that reduce binding also result in a loss of function in cells. The combination of our structure prediction, deep mutational scanning, and targeted follow-up mutations suggests a plausible MlaC-MlaA binding mode, in which the C-terminal region of MlaA likely plays a significant role in this interaction.

### AlphaFold2 prediction of MlaC-MlaD complex is consistent with mutational analysis

To better understand how MlaC interacts with MlaD, we used AlphaFold2 (Jumper et al., 2021) as implemented in ColabFold (Mirdita et al., 2022) to predict the structure of the complex. The MlaC-MlaD interaction is more complicated to model than the MlaC-MlaA interaction, because MlaD forms a homohexameric ring, and between one and six MlaC molecules may bind. We first predicted a model of a single MlaC bound to an MlaD hexamer (1 MlaC:6 MlaD) (**Fig. 4a, 4b, S4a, S4b, and S5**). The predicted structures of MlaC and MlaD have strong confidence statistics and are similar to previously solved X-ray and cryo-EM structures (Chi et al., 2020; Coudray et al., 2020; Ekiert et al., 2017; Tang et al., 2021) (**Fig. S4b and S4c**). The MlaC-MlaD interface is globally similar in the top five predictions (**Fig. S4a**), but we also observed some differences. The predicted contacts between the cleft on MlaC and a flexible loop near the periphery of the MlaD ring (β6-β7 loop) are similar across the top five models (**Fig. S4a**). However, the angle of MlaC relative to the MlaD ring varies, suggesting that this interface may function as a flexible hinge. At one extreme, MlaC is rotated outward, away from the lipid transport tunnel formed at the center of the MlaD ring (“outward” prediction), and makes few predicted contacts outside of the cleft-loop interface (**Fig. 4a and S5**). At the other extreme, MlaC is fully rotated inward, towards the MlaD tunnel (“inward” prediction), and makes additional predicted interactions with the C-terminal helices of an adjacent MlaD subunit (**Fig. 4a and 4b**). In this configuration, the entrance to the lipid binding pocket of MlaC is brought in close proximity to the MlaD tunnel, which could facilitate the transfer of lipids between MlaC and MlaD. There is extensive overlap between the footprint of MlaD and MlaA on MlaC; the same cleft on MlaC is predicted to be important for both MlaD and MlaA binding (**Fig. 4c**), and the binding of MlaD and MlaA to MlaC may be mutually exclusive. The overlapping binding interfaces may also complicate our ability to deconvolute the relative importance for MlaD vs MlaA binding of a specific residue with decreased fitness from deep mutational scanning.

**Figure 4.**
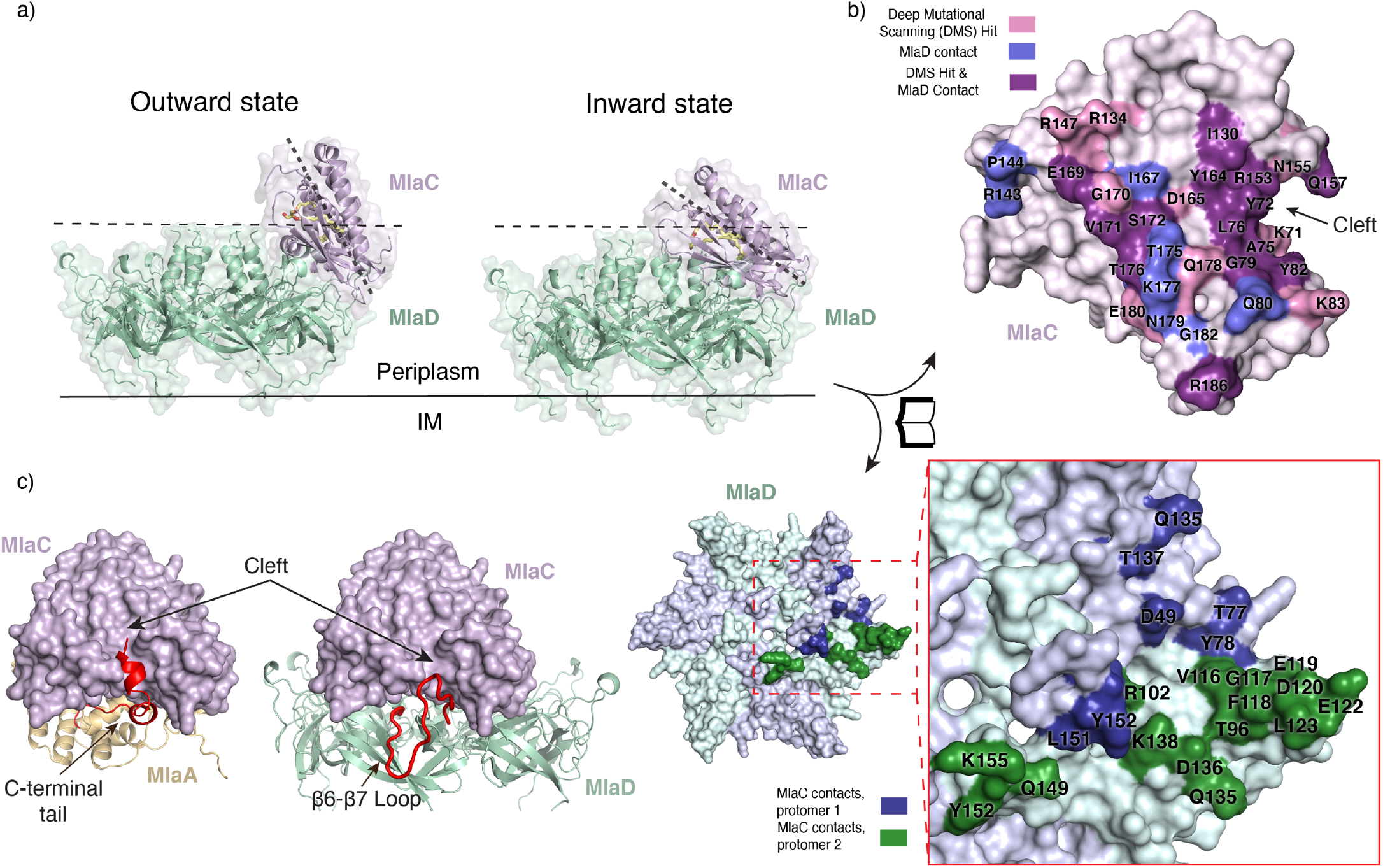
AlphaFold2 prediction of MlaC-MlaD protein-protein interface. **a)** The two extreme AlphaFold2 predictions for the MlaC-MlaD interaction. Outward state: MlaC is oriented furthest from the MlaD central pore. Inward state: MlaC is oriented closest to the MlaD central pore. Dashed lines are used to highlight the difference in binding angle of MlaC. A black line represents the IM. **b)** Open-book representation, in which MlaC and MlaD are rotated as indicated and shown as molecular surfaces, highlighting interaction interfaces in the inward state. Residues in MlaC predicted to be within 4 Å distance of MlaD from AlphaFold2, shown to have reduced fitness from deep mutational scanning (DMS), or both, are mapped onto the structure as indicated in the key. Residues in MlaD predicted to be within 4 Å distance of MlaC from AlphaFold2 are mapped onto the structure as indicated in the key. **c)** Predicted interactions of MlaC, shown as a surface, with MlaA of MlaD. The MlaA C-terminal tail (red) and the MlaD β6-β7 loop (red), are both predicted to interact with the cleft region of MlaC.

The predicted MlaD-interacting residues on MlaC are in general agreement with the results from deep mutational scanning (**Fig. 4b and S5, Supplementary Data File 1**). In the outward conformation, 12 residues on MlaC are predicted to contact MlaD, and 10 of these showed decreased fitness by deep mutational scanning (**Fig. S5, Supplementary Data File 1**). In the inward conformation, the MlaC-MlaD interface is larger. While several interactions at the cleft-loop interface are predicted in both conformations, many additional contacts are made in the inward state at a novel interface between the periplasm-facing surfaces of MlaD and the ɑ5 helix and β-sheet of MlaC. In this inward state, 25 residues on MlaC are predicted to contact MlaD, of which 14 residues were identified as important by deep mutational scanning (**Fig. 4b, Supplementary Data File 1**). Two functionally important MlaC residues identified by deep mutational scanning (A168 and R186) are predicted to interact with MlaD only in the inward state; not with MlaD in the outward state, nor with MlaA. In the absence of any other clear explanation for the functional importance of A168 and R186, decreased fitness upon mutation of these residues may indicate that MlaC samples the inward state at least some of the time.

To validate and probe further the MlaC-MlaD interface experimentally, we tested binding of 17 MlaC mutants to MlaD (**Fig. 5a, 5b and S3**). We recombinantly expressed and purified WT and mutant MlaC proteins, and tested their ability to bind immobilized MlaD protein on a sensor using biolayer interferometry. WT MlaC binds MlaD with a K_D_ of ∼18 +/-6.9 μM. Of the MlaC mutations tested, four mutants, L76D, Y82K, I130D, and R153A, showed no detectable binding to MlaD in this assay, using a maximum concentration of 100 μM MlaC (**Fig. 5a and 5b**). These mutations are clustered at the predicted cleft-loop interface of MlaC-MlaD (**Fig. 5a**). Other mutants show diminished MlaC binding to MlaD as compared to WT, including Y72A, R134E, R147D, N179E, and R186A (**Fig. S3**). Y72A is located at the cleft-loop interface, while the four other mutations are located on the MlaC β-sheet (R134E and R147D) and ɑ5 helix (N179E and R186A) which are predicted to interact with MlaD only in the inward state. These results suggest that the inward conformation may be functionally relevant.

**Figure 5.**
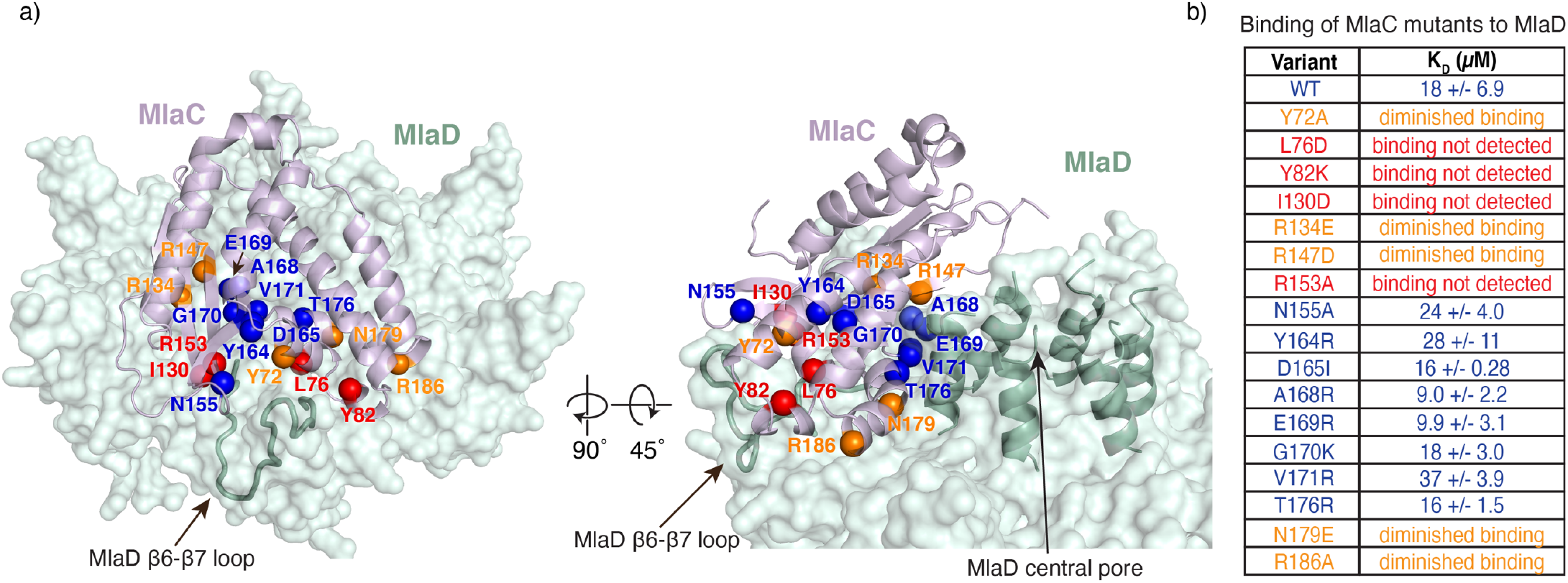
Binding experiments of MlaC mutants to MlaD probe the role of amino acids at the predicted protein-protein interface. **a)** MlaC residues predicted to be at the interaction interface with MlaD and for which mutations were generated are mapped on the predicted inward state as spheres. **b)** Summary of binding experiments for WT and mutant MlaC proteins against WT MlaD. Colors are correlated for (a) and (b): blue, binding similar to WT; orange, diminished binding compared to WT; red, no binding detected using the highest protein concentration used for WT control (100μM).

### MlaD β6-β7 loop is critical for interaction with MlaC

To probe the regions of MlaD that are important for interaction with MlaC, we first focused on the β6-β7 loop in MlaD, which is at the cleft-loop interface and makes similar interactions with MlaC in all five AlphaFold2 predictions. We generated point mutations in two hydrophobic residues in the β6-β7 loop that are predicted to interact with MlaC: F118A and L123A (**Fig. 6b**) (Ercan et al., 2019). The L123A mutation results in a ∼10-fold reduction in MlaC affinity (K_D_ of ∼200 +/-64 μM), and binding of MlaC to the F118A mutant or F118A/L123A double mutant was undetectable under our assay conditions (**Fig. 6c and S3**). These results show that point mutations in the β6-β7 loop of MlaD greatly reduce binding between MlaC and MlaD. To assess the impact of these mutations using the cell growth assay, we tested their ability to genetically complement an *E. coli mlaD* knockout strain, which is unable to grow in the presence of SDS and EDTA unless a functional copy of *mlaD* is provided on a plasmid. Surprisingly, the F118A and L123A mutations fully restored growth of an *mlaD* knockout *E. coli* strain in the presence of SDS+EDTA, perhaps indicating that the residual binding affinity of MlaC to each MlaD mutant was sufficient to support Mla function (**Fig. 6d**). Indeed, an MlaD F118A/L123A double mutant, which presumably would have even lower affinity for MlaC, was unable to complement the *mlaD* knockout. Western blotting against cell lysates from the strains used for the cell growth assay revealed consistent expression of each mutant protein, suggesting the differences in cell growth are not due to altered expression levels (**Fig. S6**). Note that MlaD mutant proteins expressed from a plasmid are highly over-expressed compared to endogenous MlaD, which in cells may overcome the effect of mutations resulting in reduced binding affinities, such as F118A and L123A. Overall, our data suggest that the interaction between MlaC and the β6-β7 loop in MlaD is a key determinant, and is required for Mla function in cells.

**Figure 6.**
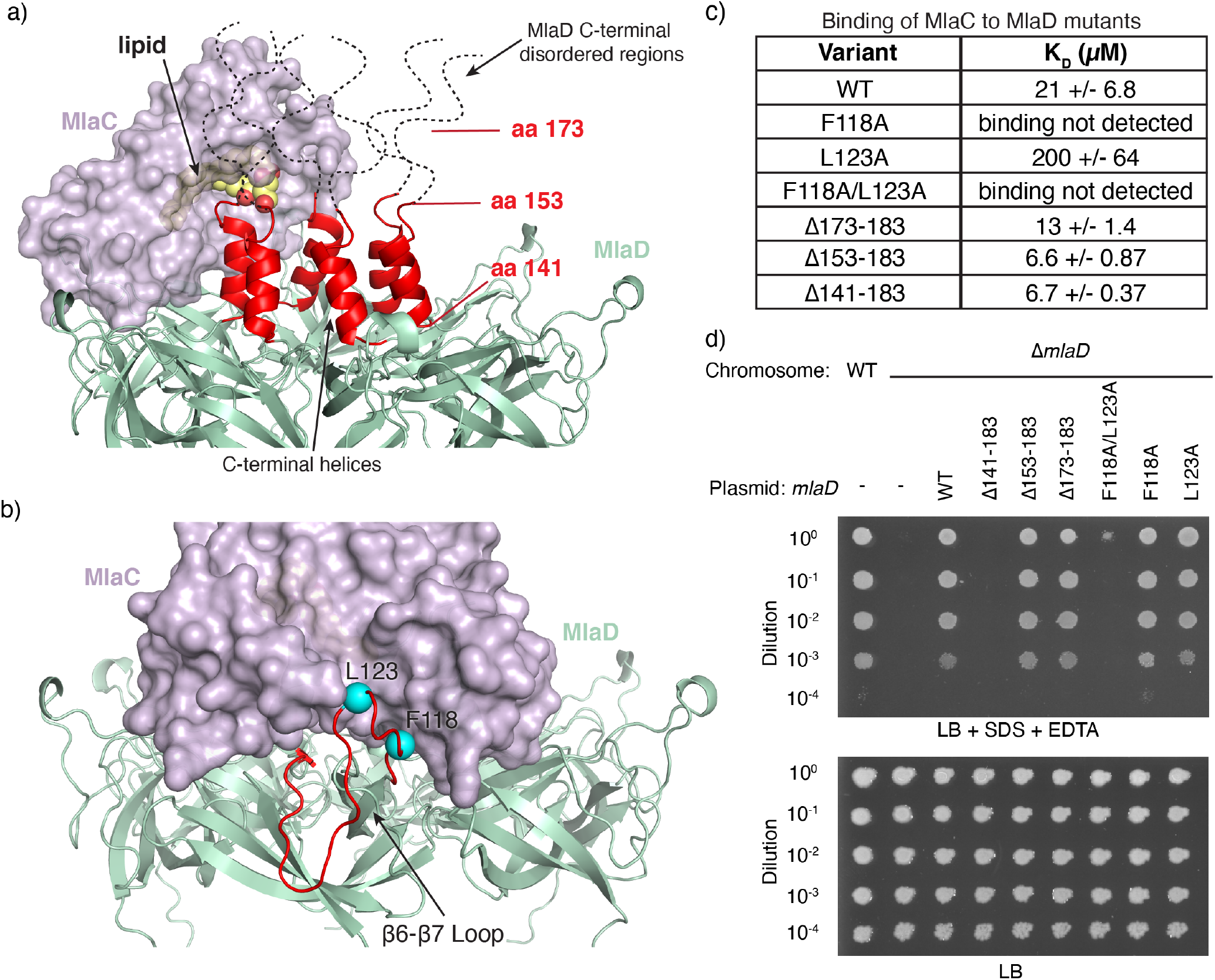
Mutations in key residues on the MlaD β6-β7 loop impair binding. **a)** The predicted inward state of MlaC (purple surface) bound to MlaD (green cartoon). MlaC is oriented close to the C-terminal helices of MlaD (red). Additional residues at the C-terminus of MlaD, which have not been resolved in any structures and are likely to be disordered, are represented by black dashed lines, drawn to scale. A lipid (yellow), not included during the MlaC-MlaD prediction, is shown based on its position in MlaC (PDB 5UWA). The orientation of MlaC positions the lipid towards the central pore of MlaD, formed by the C-terminal helices. Positions for MlaD C-terminal truncation mutants are shown in red with lines indicating the boundary of the deletion. **b)** The predicted inward state of MlaC-MlaD. The MlaD β6-β7 loop is red, and two residues predicted to interact, F118 and L123, are colored in cyan. **c)** Summary of binding experiments for MlaC against WT and mutant MlaD proteins. **d)** Genetic complementation of Δ*mlaD E. coli* cells with plasmids expressing WT MlaD, point mutations, or C-terminal truncations. 10-fold serial dilutions of strains were spotted on LB agar with or without SDS+EDTA as indicated. All plates contain 2% arabinose to induce *mlaD* expression from plasmid.

To assess the importance of interactions between MlaC and the C-terminal helices and disordered tails of MlaD (Coudray et al., 2020; Ekiert et al., 2017) (**Fig. 6a**), we constructed three sequential truncation mutants: MlaDΔ173-183 and MlaDΔ153-183, which delete disordered regions, and MlaDΔ141-183, which deletes the disordered regions and C-terminal helices. We found that MlaC binds to all three truncation mutants with affinities similar to WT MlaD, suggesting that interactions between MlaC and the C-terminal helices and disordered tails of MlaD are not critical for binding (**Fig. 6c and S3**). In cell growth assays, the MlaDΔ173-183 and MlaDΔ153-183 mutants fully rescue the growth of an *E. coli mlaD* knockout strain in the presence of SDS and EDTA (**Fig. 6d**). However, the MlaDΔ141-183 mutant does not rescue growth, as previously reported (Ekiert et al., 2017). Western blotting against cell lysates from the strains used for the cell growth assay revealed consistent expression of each mutant protein (**Fig. S6**). Taken together, these data suggest that the C-terminal helices and disordered regions of MlaD are dispensable for MlaC binding. The C-terminal helices of MlaD appear to be important for function in our cell growth assay, perhaps due to a role in lipid transfer or a structural role as part of the lipid transport tunnel through the MlaD ring.

### Cryo-EM reveals compositional heterogeneity of MlaC binding to MlaFEDB

To gain additional insights into MlaC binding to MlaFEDB, we turned to single particle cryo-EM. Initial attempts at preparing a stable MlaC-MlaFEDB complex were unsuccessful, likely because the affinity of MlaC for MlaD is very low. To increase the affinity of MlaC for MlaD and stabilize the MlaC-MlaFEDB complex, we designed an artificially trimerized construct by tethering MlaC to the foldon trimerization domain of T4 fibritin via long Ser-Gly linkers (Ekiert et al., 2009; Meier et al., 2004; Tao et al., 1997) (Supplementary Table 3). This construct binds MlaD with ∼1000-fold higher affinity (apparent K_D_ = 21.9 +/-6.0 nM), likely through increased avidity due to multivalent binding (**Fig. S3**). We mixed trimerized MlaC and MlaFEDB complex at a 1:1 molar ratio and collected cryo-EM data for this sample. The data revealed a large amount of compositional and conformational heterogeneity (**Fig. S7**), which limits our current analysis to low resolution. Based on previous high-resolution structures of MlaFEDB (Chi et al., 2020; Coudray et al., 2020; Mann et al., 2021; Tang et al., 2021; Zhang et al., 2020; Zhou et al., 2021), we can unambiguously assign most of the density from our reconstruction to the MlaFEDB complex. After accounting for MlaFEDB, additional density is observed adjacent to the MlaD ring, consistent with the size and shape of MlaC (**Fig. 7a**). 3D classification revealed several maps differing in the relative locations of the MlaC molecules bound around the MlaD hexamer (**Fig. S7**). In most reconstructions, we see two clear densities that we interpret to be MlaC, based on their size and shape. In all observed configurations, MlaC molecules do not bind adjacent MlaD protomers, leaving 1-2 binding sites vacant in between. This suggests that MlaC binding to adjacent MlaD protomers is disfavored, perhaps due to steric clashes between neighboring MlaC molecules. To investigate this hypothesis further, we predicted a complex of six MlaC molecules bound to a hexameric MlaD ring using AlphaFold2, which yielded 5 nearly identical models. This predicted 6:6 complex suggests that, while 6 MlaCs can physically be accommodated on an MlaD hexamer in an “outward-like” state, it would require very tight lateral packing between adjacent molecules, and would likely be unfavorable (**Fig. S4d and S4e**). Thus, 2-3 MlaC molecules can likely bind to one MlaFEDB complex *in vitro*, though it remains unclear if more than one MlaC can simultaneously bind an MlaFEDB complex in cells.

**Figure 7.**
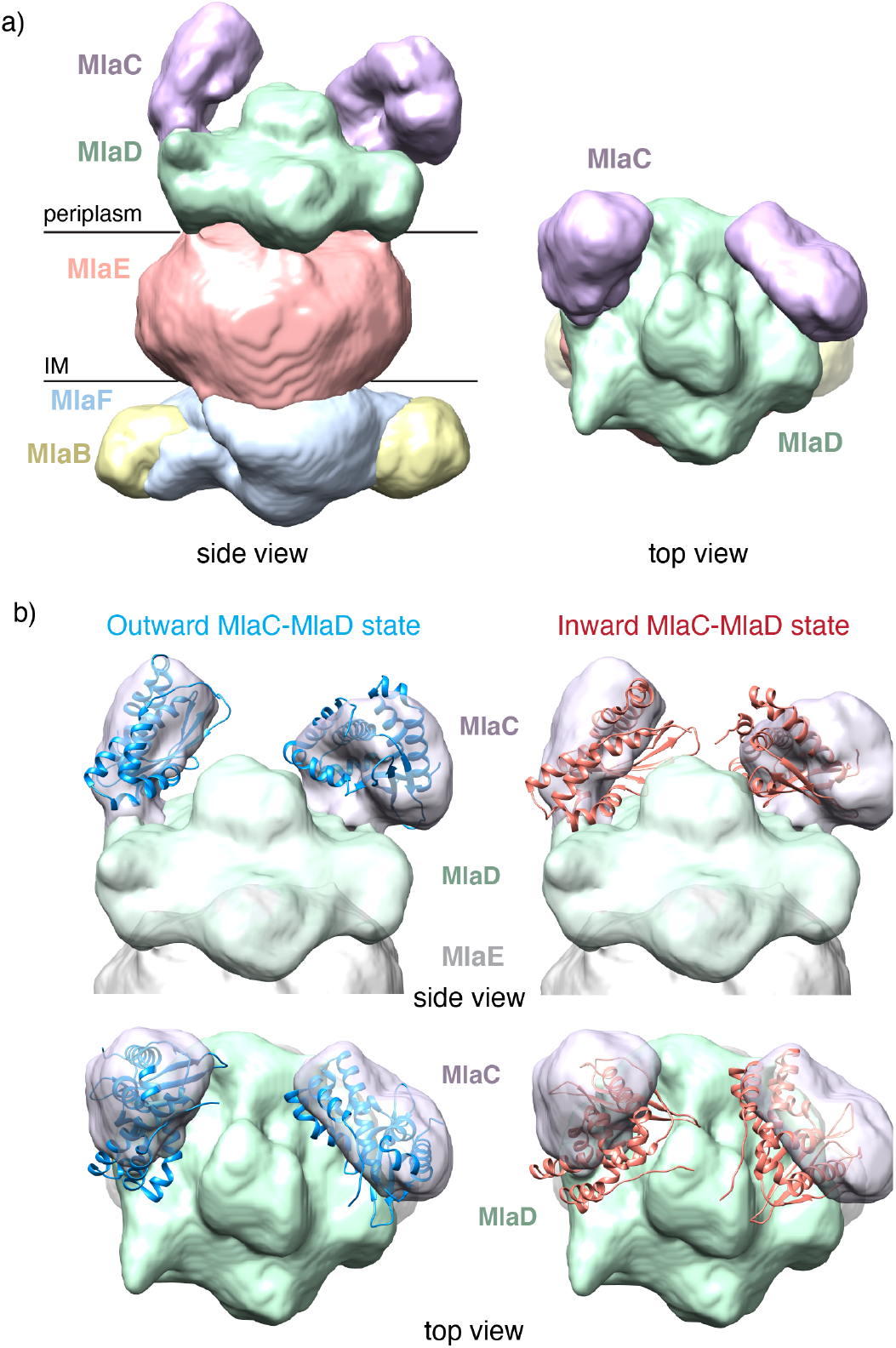
Low resolution cryo-EM maps reveal two MlaC molecules can bind to MlaD in agreement with outward state prediction. **a)** Cryo-EM reconstruction (class 1) shows two additional densities (purple) atop the MlaFEDB complex, in proximity to MlaD. Purple, densities presumed to be MlaC; green, MlaD; pink, MlaE with detergent micelle; blue, MlaF; yellow, MlaB. High-resolution structures for MlaFEDB are available (PDB 6XBD), allowing us to unambiguously assign density corresponding to the MlaFEDB complex, and analyze additional density, presumably corresponding to MlaC. **b)** The predicted inward (blue) and outward (red) states of the MlaC-MlaD complex were fit into the density (class 1) as rigid bodies, and MlaC in each of these states is shown in cartoon representation.

To better understand the conformation of MlaC bound to MlaD, we fit the AlphaFold2 predictions of the inward and outward states as rigid bodies into our EM density maps. We found that in all maps, the outward state fits the experimental density better than the inward state (**Fig. 7b**), suggesting that under the conditions of our cryo-EM experiment, the outward state is predominant. Thus, if MlaC indeed samples the inward state, it may be transient and coupled to lipid transfer, or dependent upon the lipid binding state of MlaC (Hughes et al., 2019) or the conformation of the rest of the MlaFEDB complex.

## Discussion

The mechanism of how lipids are transferred between MlaC and the MlaFEDB complex, and between MlaC and the MlaA-OmpC/F complex have remained unclear. This is due in part to the underlying question of how MlaC interacts with the two complexes. To address this question, we used a combination of deep mutational scanning, AlphaFold2 structure prediction coupled with low resolution EM maps, binding experiments and cell growth assays. The deep mutational scanning provided an unbiased insight into functionally important amino acids in MlaC. The roles of several residues identified as being important can be attributed to known functions such as the signal peptide and the lipid binding pocket (Ekiert et al., 2017; Hughes et al., 2019). Other regions on MlaC where mutations had high fitness costs were surface-exposed, and we propose a role for these in MlaD and MlaA binding. Our results suggest that the residues with the greatest impact on MlaD and MlaA interactions are localized to the MlaC cleft, and suggest that MlaC can only bind to MlaA or MlaD, but not both simultaneously. Of the 25 solvent exposed residues from deep mutational scanning, only two residues (E63 and D165) are not predicted to interact with either MlaD or MlaA. It may be possible that these residues are involved in other protein functions such as lipid transfer or may affect protein folding or stability.

These data together with previous work lead to a possible model for protein-protein interactions and lipid transfer between the key components of the Mla pathway (**Fig. 8**). At the OM, MlaC interacts with MlaA such that a lipid bound in the MlaA channel is poised for transfer to the lipid-binding pocket of MlaC. Because MlaC binds lipids with very high affinity, it has previously been suggested that lipid transfer from MlaA to MlaC may be energetically favorable and occur spontaneously (Abellón-Ruiz et al., 2017; Ercan et al., 2019). The C-terminal tail of MlaA is an important part of the MlaC-MlaA interaction, though other parts of the interface, such as the ɑ6 helix, may also play a role in binding and lipid transfer. In crystal structures of free MlaA-OmpF/C complexes, electron density is not observed for the MlaA C-terminal tail, suggesting that it is unstructured or flexible in the absence of MlaC but undergoes a disorder-to-order transition upon MlaC binding. After ferrying the bound phospholipid across the periplasmic space, MlaC binds to the MlaFEDB complex at the inner membrane via an interaction between the MlaC cleft and the β6-β7 loop of MlaD. Based on our cryo-EM analysis, it is possible that 1, 2, or perhaps 3 MlaC proteins may interact with MlaD simultaneously. The interaction between MlaC and MlaD is likely dynamic around a hinge point that results in MlaC being further away from the MlaD central pore or closer to it. The different conformations of MlaC relative to MlaD may play a role in lipid transfer between the two proteins, by bringing the lipid-binding pocket of MlaC in close proximity to the lipid transport tunnel through the MlaD ring. Sampling of the inward state is supported by mutagenesis data presented here, as well as previous work suggesting that residues close together in the inward state but distant in the outward state can be crosslinked *in vivo* (Ercan et al., 2019), whereas cryo-EM data are more consistent with the outward state. Many open questions remain with respect to the Mla mechanism and lipid transfer between MlaC and MlaD. Structural insights at higher resolution will be necessary to understand this process, and how conformational changes may be transmitted in an ATP-dependent manner to promote lipid release from MlaC and transfer to MlaFEDB.

**Figure 8.**
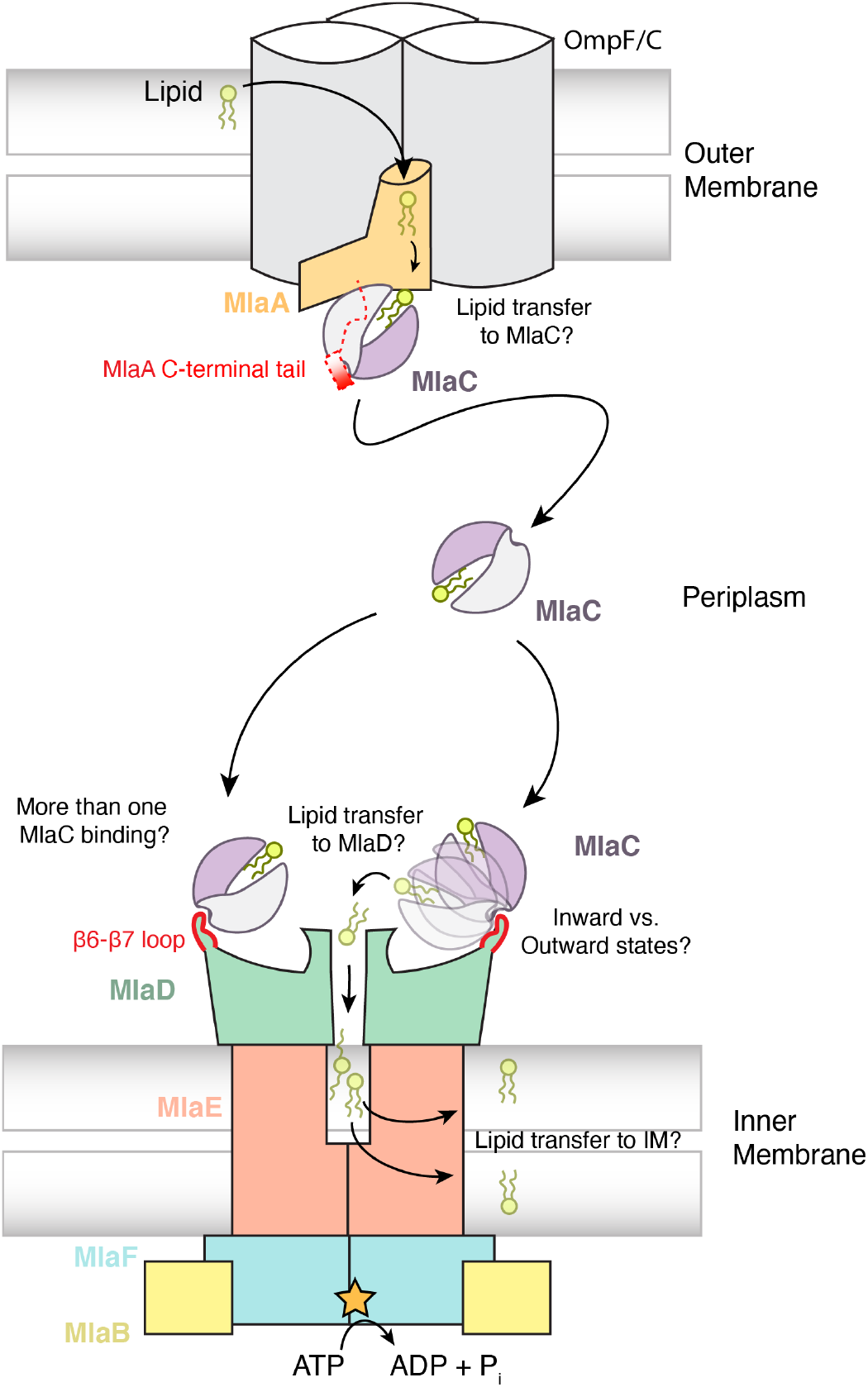
A model for MlaC interaction with MlaD and MlaA for lipid transport, in the context of import. MlaC represents a clamshell, in which one half (grey) predominantly makes interactions with both binding partners, MlaA and MlaD. At the OM, MlaC interacts with MlaA, and a major component of this interaction is the MlaA C-terminal tail, which binds the cleft of MlaC. Predicted interactions of MlaC with MlaA suggest that the MlaC lipid-binding pocket is poised for lipid transfer from MlaA upon binding. Upon disassociation from MlaA, MlaC traverses the periplasm and interacts with MlaD in the IM MlaFEDB ABC transporter complex. Two MlaC molecules may simultaneously bind at the outer loops of MlaD and this interaction may be flexible around a hinge-point, resulting in inward or outward-bound states of MlaC to MlaD. In the inward state, MlaC is oriented closer to the MlaD central pore, possibly facilitating lipid transfer. Lipid transfer and release is facilitated by conformational changes in MlaFEDB, driven by ATP hydrolysis.

## Methods

### MlaC deep mutational scanning

The complete single-mutation library of MlaC was constructed by oligonucleotide-directed mutagenesis, using an NNS codon encoding (where “N” represents any base, and “S” represents either G or C) to randomize each site in the open reading frame. To ensure near full coverage of the MlaC deep mutational scanning library by paired-end sequencing (2×250), we split the MlaC sequence into two groups (AA positions 1-107 and 108-211). The two groups were kept as distinct libraries throughout the cloning process and experiment. NNS libraries for each codon were amplified separately in a two-step fusion PCR reaction, and the resulting PCR products were mixed in an equimolar ratio. The pooled NNS MlaC library PCR fragments and a pBAD-derived arabinose inducible vector (pBEL2403) were digested with the restriction enzyme, SfiI, followed by ligation with T4 ligase. TOP10 *E. coli* cells were transformed with the ligated product, and plasmids were purified with the Zymo miniprep kit to yield the “pre-selection” plasmid library. Next, the Δ*mlaC* strain, bBEL464, was transformed with the complete NNS library (AA positions 1-107 or 108-211) and grown overnight on LB agar plates with carbenicillin. The colonies were then scraped and pooled, followed by plating on either LB + 2% arabinose (“no selection”) or LB + 2% arabinose + 1.3 mM EDTA + 1% SDS (“selection”). Lastly, after overnight growth on the no selection and selection plates, colonies from each condition were separately scraped and pooled, plasmids were extracted, and amplicons were generated by PCR for amplicon sequencing by Illumina 2×300 paired-end sequencing. We performed two independent biological replicates of the deep mutational scanning experiment starting from the same libraries (**Fig. S1a and S1b**).

Paired-end sequencing data was first processed by removal of sequencing adaptors (Trimmoatic v0.39). Next, reads were mapped to a reference WT MlaC sequence using the bowtie2 algorithm (v2.4.1), filtered with samtools (v1.9; flags -f 2 -q 42), and overlapping paired ends were merged into a single sequence with pandaseq (v2.11). Lastly, primer sequences used for amplicon amplification were removed using cutadapt (v1.9.1). Processed and merged reads were then analyzed using custom python scripts to count the frequency of the MlaC variants (**Fig. S1c**). Briefly, DNA sequences were filtered by length, removing any sequence larger or smaller than the length of the expected library. Next, sequences were correctly oriented to the proper reading frame and translated to the corresponding protein sequence. Finally, the frequency of each amino acid variant at every position was counted and the counts were normalized to the sequencing depth as read counts per million. These normalized counts were then used for calculation of the mutational cost, which is defined as the log frequency of observing each amino acid *x* at each position *i* in the selected versus the non-selected population, relative to the wild type amino acid (McLaughlin et al., 2012). The equation for this calculation is as follows:

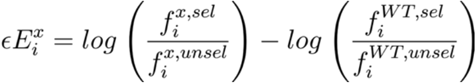

### AlphaFold2 predictions

In generating structural predictions of the interactions between MlaA and MlaD with MlaC, we utilized several AlphaFold prediction softwares, including AlphaFold2, AlphaFold Multimer via Cosmic2 (Cianfrocco et al., 2017; Evans et al., 2022; Jumper et al., 2021), and AlphaFold2_advanced via Colabfold. Sequences of MlaC, MlaD, and MlaA from *E. coli* were retrieved from the MG1655 reference genome in the NCBI database. In predictions with MlaC, the signal sequence (amino acids 1-21) was excluded. In predictions with MlaD, the transmembrane region (amino acids 1-31) and disordered C-terminal region (amino acids 156-183) of the protein were excluded. In predictions with MlaA, the signal sequence (amino acids 1-27) was excluded. For the MlaC-MlaA complex, we used AlphaFold-Multimer to predict a 1:1 complex. For the MlaC-MlaD complex, we used AlphaFold2 to predict a 6:6 complex and AlphaFold2_advanced to predict a 1:6 complex. Default program parameters were used in predictions. Program outputs yielded 5 ranked models of protein-protein predicted interaction. Predicted alignment error (PAE) plots were generated using AlphaPickle (Arnold, M. J. (2021) AlphaPickle doi.org/10.5281/zenodo.5708709).

### Phenotypic assays for *mla* mutants in *E. coli*

Knockouts of *mlaA, mlaD* and *mlaC* were constructed in *E. coli* BW25113 by P1 transduction from corresponding strains of the Keio collection (**Supplementary Table 4**) (Baba et al., 2006), followed by excision of the antibiotic resistance cassettes using pCP20 (Cherepanov and Wackernagel, 1995). To test the impact of the various *mlaA, mlaD* and *mlaC* mutants on protein function in cells, pBAD-derived expression plasmids encoding the mutant of interest were transformed into the appropriate knockout strain and sensitivity to SDS+EDTA was tested essentially as previously described (Kolich et al., 2020). For MlaA and MlaC mutants, serial dilutions of overnight cultures were performed in 96-well plates and then spotted with a micropipet or pin replicator (“frogger”) on plates containing LB agar (BD Difco #244510) and 2% arabinose (CHEM-IPEX #01654) (control plate) or LB agar supplemented with 2% arabinose, 0.25% SDS (Sigma L5750) and 0.35-0.40mM EDTA (Sigma ED2SS) (selection plate) for *mlaA* and *mlaC*. For MlaD mutants, dilutions were spotted on LB agar plates without arabinose, as the leaky expression from the uninduced plasmids was sufficient for complementation. The plates were then incubated for 16h at 37°C.

For experiments involving *mlaA* mutations, we introduced the mutation of interest into plasmid pBEL2511, which expresses WT MlaA under the control of an arabinose-inducible promoter. The following mutations were tested: Δ227-251 (pBEL2513), Δ238-251 (pBEL2512), Δ244-251 (pBEL2665), Ile241Asn (pBEL2639), Leu245Asn (pBEL2640), Ile248Asn (pBEL2641), Phe223Asn (pBEL2663), Leu230Asn (pBEL2664) and Asp198Asn (pBEL2644).

For experiments involving *mlaC* mutations, we introduced the mutation of interest into plasmid pBEL2403, which expresses WT MlaC under the control of an arabinose-inducible promoter. The following mutations were tested: Ala34Arg (pBEL2521), Val68Arg (pBEL2541), Tyr105Ala (pBEL2547), Ala108Arg (pBEL2522), Gln115Ala (pBEL2534), Val135Arg (pBEL2542), Ile137Asp (pBEL2530), Val146Arg (pBEL2543), Ala163Arg (pBEL2523), Ala168Arg (pBEL2524), Glu169Arg (pBEL2526), Tyr72Ala (pBEL2545), Leu76Asp (pBEL2531), Tyr82Lys (pBEL2546), Ile130Asp (pBEL2528), Arg134Glu (pBEL2535), Arg147Asp (pBEL2536), Arg153Ala (pBEL2537), Asn155Ala (pBEL2532), Tyr164Arg (pBEL2548), Asp165Ile (pBEL2535), Gly170Lys (pBEL2527), Val171Arg (pBEL2544), Thr176Arg (pBEL2540), Asn179Glu (pBEL2533) and Arg186Ala (pBEL2538).

For experiments involving *mlaD* mutations, we introduced the mutation of interest into plasmid pBEL1195, which expresses the complete MlaFEDCB operon under the control of an arabinose-inducible promoter. The following mutations were tested: Δ141-183 (pBEL1330), Δ153-183 (pBEL1936), Δ173-183 (pBEL1935), Phe118Ala/Leu123Ala (pBEL1953), Phe118Ala (pBEL1933) and Leu123Ala (pBEL1934). Further plasmid details can be found in **Supplementary Table 3**.

### Western blot to detect MlaA, MlaD and MlaC expression level from complementation assay

Western blots were performed in order to assess the expression level of MlaA, MlaD and MlaC WT and their mutants on complementation assays plates. The samples were prepared by scraping a zero dilution spot of each sample from the control plates and resuspended in 1mL 1XPBS. The OD of each sample was normalized to 0.25 and 1mL of the cells were centrifuged at 5000 g, at 4°C for 30 min. After removing the supernatants from the centrifugation step, the pellets were resuspended in 50 μL of 1x SDS sample loading buffer. The samples were boiled at 95°C for 10 min to break open the cells, then 15μL of each sample was loaded and run on SDS-PAGE gels at 120V. Proteins in the gels were transferred onto nitrocellulose membrane from Biorad (#L002050A) using Turbo transfer buffer and a TransBlot Turbo from Biorad (#10026938). After transfer, the membranes were washed 2 times for 10 minutes with PBST buffer (10% PBS v/v and 0.05% of Tween 20) before blocking with PBST buffer containing 5% BSA, at room temperature for 1 h. The membranes were then washed 3 times with PBST buffer and incubated with primary antibody buffer (PBST buffer containing 5% of BSA, 0.02% Sodium Azide), anti-mouse GAPDH antibody at 1μg/mL from Abcam #ab125247, and the respective antibody: anti-rabbit MlaC antibody (0.8μg/mL from Capra Science #1257.1286) and MlaD antibody (provided by Henderson lab, University of Queensland) for 1 h at room temperature. The membranes were washed 3 times with PBST buffer for 10 minutes and incubated with secondary antibodies (PBST buffer + 5% BSA + 0.1μg/mL of both goat anti-rabbit 800nm (LI-COR Biosciences; #925-32211) dilution 1:10,000 and goat anti-mouse 680nm antibodies dilution 1:10,000 (LI-COR Biosciences; #926-68070)). Finally, the membranes were washed 3 times for 10 min with PBST buffer before imaging using the LI-COR Odyssey XF (LI-COR Biosciences).

### Expression and purification of soluble constructs

The following soluble constructs were used, as described in Supplementary table 3.

- WT MlaC (pBEL 1203)
- MlaC mutants: Ala168Arg (pBEL2552), Asp165Ile (pBEL2553), Glu169Arg (pBEL2554), Gly170Lys (pBEL2555), Ile130Asp (pBEL2556), Leu76Asp (pBEL2559), Asn155Ala (pBEL2560), Asn179Glu (pBEL2561), Arg134Glu (pBEL2563), Arg147Asp (pBEL2564), Arg153Ala (pBEL2565), Arg186Ala (pBEL2566), Thr176Arg (pBEL2568), Val171Arg (pBEL2572), Tyr72Ala (pBEL2573), Tyr82Lys (pBEL2574), Tyr164Arg (pBEL2576)
- MlaC-foldon (pBEL2319)
- WT MlaD (pBEL1160)
- MlaD mutants: Δ141-183 (pBEL1161), Δ153-183 (pBEL1224), Δ173-183 (pBEL1223), Phe118Ala (pBEL1909), Leu123Ala (pBEL1910), Phe118Ala/Leu123Ala (pBEL1954) WT sfGFP (pBEL1092)
- sfGFP chimera mutants: sfGFP-MlaA227-251 (pBEL2707), sfGFP-MlaA 227-251/I241N (pBEL2753), sfGFP-MlaA227-251/L245N (pBEL2754), sfGFP-MlaA227-251/I248N (pBEL2755), MlaA (227-237) (pBEL2756), MlaA (227-243) (pBEL2757)

The expression plasmid was transformed into Rosetta 2 (DE3) cells (Novagen). 20 mL LB with 38 μg/mL chloramphenicol and 100 μg/mL carbenicillin was inoculated with a single Rosetta 2 (DE3) colony containing the plasmid of interest. The culture was grown for 16 h at 37 °C, shaking at 200 rpm. Overnight cultures of Rosetta 2 (DE3) containing the relevant expression plasmid were diluted 1:50 in LB (Difco) supplemented with carbenicillin (100 μg/mL) and chloramphenicol (38 μg/mL) and grown at 37°C with shaking at 200 rpm. Cultures were induced at OD_600_ = ∼0.6 by adding IPTG to a final concentration of 1 mM, and induction was allowed to proceed for 4 hr. The cultures were then harvested by centrifugation at 5,000 g for 30 min at 4°C, and cell pellets were resuspended in lysis buffer (50mM Tris pH 8.0, 300mM NaCl). The resuspended cells were flash-frozen in liquid nitrogen and stored at −80°C. Cells were thawed at room temperature and lysed by passing three times through an Emulsiflex-C3 cell disruptor (Avenstin). Lysate was centrifuged at 35,000 g for 30 min to pellet cell debris. The supernatant was then passed through 1.5 mL bed volume of NiNTA resin (GE Healthcare #17531802) twice, and washed with ∼130 column volumes of Ni Wash Buffer (50 mM Tris pH 8.0, 300 mM NaCl, 40 mM imidazole) to remove non-specifically bound proteins. Target proteins were eluted with 15 mL Ni Elution Buffer (50 mM Tris pH 8.0, 300 mM NaCl, 250 mM imidazole). Eluted protein was pooled and concentrated to 0.5 mL. Size exclusion chromatography was performed on the concentrated protein using a Superdex 200 Increase 10/300 gel filtration column (GE Healthcare) equilibrated in gel filtration buffer (10 mM Na_2_HPO_4_.7H_2_O, 137 mM NaCl, 2.7 mM KCl, 1.5 mM KH_2_PO_4_). The peak containing our protein of interest was pooled, concentrated, and stored on ice at 4°C for use in biochemical binding experiments, or aliquoted and flash frozen in liquid nitrogen and stored at −80°C for future use.

### Binding experiments via Biolayer Interferometry

Biolayer interferometry using the Octet Red96 instrument (ForteBio) was used to measure binding interactions of MlaC to WT and mutant MlaD proteins, as well as WT and mutant GFP. Proteins to be immobilized on octet sensors (also called the “load” proteins) were non-specifically biotinylated. This includes the following constructs, as described in Supplementary Table 3:

- WT MlaD (pBEL1160)
- MlaD mutants: Δ141-183 (pBEL1161), Δ153-183 (pBEL1224), Δ173-183 (pBEL1223), Phe118Ala (pBEL1909), Leu123Ala (pBEL1910), Phe118Ala/Leu123Ala (pBEL1954) WT sfGFP (pBEL1092)
- sfGFP chimera mutants: sfGFP-MlaA227-251 (pBEL2707), sfGFP-MlaA 227-251/I241N (pBEL2753), sfGFP-MlaA227-251/L245N (pBEL2754), sfGFP-MlaA227-251/I248N (pBEL2755), sfGFP-MlaA227-237 (pBEL2756), sfGFP-MlaA227-243 (pBEL2757)

For non-specific biotinylation, the purified and concentrated protein was incubated with 1:3 protein to biotin ratio of 1mM NHS-PEG4-Biotin (VWR #PI21362) for 1 hr at room temperature. The biotinylated protein was separated from excess biotin on a Superdex 200 Increase 10/300 gel filtration column (GE Healthcare) equilibrated in 1x PBS buffer (10 mM Na_2_HPO_4_·7H_2_O, 137 mM NaCl, 2.7 mM KCl, 1.5 mM KH_2_PO_4_). The peak corresponding to biotinylated protein was pooled, concentrated to 30 mg/mL, flash frozen in liquid nitrogen and stored at - 80°C. Streptavidin (SA) biosensors (ForteBio) were used to immobilize biotinylated load protein on the sensor. Prior to loading the protein, sensors were pre-incubated with freshly made 1x Kinetics buffer (0.01 mg/mL BSA, 0.02 % Tween 20, 10 mM Na_2_HPO_4_ ·7H_2_O, 137 mM NaCl, 2.7 mM KCl, and 1.5 mM KH_2_PO_4_) for 10 min at 30°C. The following assay was used for binding experiments: Baseline step, in buffer only, 60 s; Load step, in which biotinylated protein was loaded onto the sensor, 5 min; Wash step, in 1x Kinetics to remove unbound load protein, 100 s; Association step in which several concentrations (with a maximum analyte concentration varying from 30 μM to 100 μM) of MlaC protein (analyte) in solution were allowed to associate with to the load protein, 30 s; Dissociation step, in which the sensor containing load protein and bound analyte is moved into 1x Kinetics buffer to remove analyte, 100 s; Equilibration step in which the sensor with load protein bound is moved into a fresh well containing 1x Kinetics buffer to stabilize signal before the following association step. A no-load control was used to confirm that the analyte did not bind non-specifically to the sensor. All steps were performed with the plate shaking at 1000 rpm and a temperature of 30°C. For all binding experiments with fast dissociation rates that converge to baseline, a single sensor was used for the entire experiment to minimize sensor-to-sensor variation. The data were processed and fit globally grouped by sensor, using a 1:1 binding model in the ForteBio analysis software (v9.0). The kinetic constant (*K*_*D*_) was calculated from curve fitting post-analysis. The average *K*_*D*_ and standard deviation is reported from at least 2 biological replicates of freshly purified analyte samples.

### Expression and purification of MlaFEDB for Cryo-EM

Plasmid pBEL1200 (Ekiert et al., 2017), which contains the *mlaFEDCB* operon with an N-terminal His-tag on MlaD, was transformed into Rosetta 2 (DE3) cells (Novagen). 20 mL LB with 38 μg/mL chloramphenicol and 100 μg/mL carbenicillin was inoculated with a single Rosetta 2 (DE3) colony containing the plasmid of interest. The culture was grown for 16 h at 37 °C, shaking at 200 rpm. For expression, overnight cultures of Rosetta 2 (DE3)/pBEL1200 were diluted 1:100 in LB (Difco) supplemented with carbenicillin (100 μg/mL) and chloramphenicol (38 μg/mL) and grown at 37°C with shaking at 200 rpm. Cultures were induced at OD_600_ = ∼0.6 by adding L-arabinose to a final concentration of 0.2%, and induction was allowed to proceed for 4 hr at 37°C. Cultures were harvested by centrifugation at 5,000 g for 30 min, and the pellets were resuspended in lysis buffer (50 mM Tris pH 8.0, 300 mM NaCl, 10% glycerol). Cells were thawed and lysed by passing three times through an Emulsiflex-C3 cell disruptor (Avenstin). Lysate was centrifuged at 15,000 g for 30 min to pellet cell debris. The clarified lysates were ultracentrifuged at 37,000 rpm (F37L Fixed-Angle Rotor, Thermo-Fisher) for 45 min to isolate membranes. Membranes were resuspended in membrane solubilization buffer (50 mM Tris pH 8.0, 300 mM NaCl, 10% glycerol, 25 mM DDM) using a paint brush and incubated overnight with gentle rocking at 4°C. The solubilized membranes were ultracentrifuged at 37,000 rpm (F37L Fixed-Angle Rotor, Thermo-Fisher) for 45 min, to pellet any insoluble material. The supernatant was incubated with 1 mL bed volume NiNTA resin (GE Healthcare #17531802) at 4°C for 60 min. The resin was washed with 200 column volumes of Ni Wash Buffer (50 mM Tris pH 8.0, 300 mM NaCl, 40 mM imidazole, 10% glycerol, 0.5 mM DDM) and bound proteins eluted with 15 mL Ni Elution Buffer (50 mM Tris pH 8.0, 300 mM NaCl, 250 mM imidazole, 10% glycerol, 0.5 mM DDM). MlaFEDB-containing fractions eluted from the NiNTA column were pooled and concentrated to 0.5 mL before separation on a Superdex 200 Increase 10/300 gel filtration column (GE Healthcare) equilibrated in gel filtration buffer (10 mM Na_2_HPO_4_·7H_2_O, 137 mM NaCl, 2.7 mM KCl, 1.5 mM KH_2_PO_4_, 0.5 mM DDM). Fractions from the peak corresponding to the MlaFEDB complex were pooled and concentrated to 8 mg/mL for cryo-EM sample preparation.

### Cryo-EM grid preparation and data collection

Both the MlaFEDB sample in 0.5mM DDM and 1x PBS and the MlaC-foldon sample in 1x PBS were thawed on ice, combined in a 1:1 molar ratio (0.2 mg/mL MlaFEDB and 0.5 mg/mL MlaC-foldon), and incubated together at room temperature for 20 s. 2.5 μL of the combined sample (at a final concentration of 0.2 mg/mL) was applied to a 2/2 300 mesh Quantifoil continuous carbon grid, which had been glow discharged for 5 s. The sample was blotted for 2.0 seconds and plunge-frozen in liquid ethane using the FEI Vitrobot Mark IV. Grid screening was performed on a Talos Arctica TEM equipped with a Gatan K3 camera at 0.548 Å/pixel, operated at 200 kV, and located at NYU School of Medicine. 3,449 un-tilted and 3,875 tilted movies were acquired via Leginon (Cheng et al., 2021; Suloway et al., 2005) on an Arctica microscope at 200 kV equipped with a Gatan K3 camera, at 0.548 Å/pixel in super-resolution mode with parameters as in Supplementary Table 1.

### Cryo-EM data processing

The data processing workflow is summarized in **Supplementary Figure 7**. Movies were aligned using MotionCor2 (Zheng et al., 2017) under control of Appion and dose weighted according to the dose measured by Leginon (Cheng et al., 2021). CryoSparc v3 (Punjani et al., 2017) was used for all following steps. Particle picking and classification strategies were optimized using a subset of 350 micrographs from which ab-initio models were also generated. Particles from un-tilted and tilted micrographs were picked using blob picker, and followed by topaz (Bepler et al., 2019) for tilted micrographs. After several rounds of 2D classification, further cleaning was done running 3D heterogeneous refinement jobs using the ab-initio and decoy reconstructions as references. 3D variability analysis was also performed and the first components of the 3D variability analysis revealed maps with one or two additional densities at a location where MlaC is predicted to bind to MlaD in the MlaFEDB complex. Several approaches were tested to tease apart the different conformations (for example, 3D variability in cluster mode, 3D classification without alignment without a mask, and with various masks). The strategy that yielded the most informative results incorporated heterogeneous refinement using frames from the 3D variability analysis as seeds. Further removal of junk was done via 2D classification, and non-uniform refinement of each class led to maps with the best local resolution around the MlaD/MlaC domains, showing either one or two MlaC connected to different monomers of MlaD. Although the resolution is low, we were able to use the cryo-EM density from several classes for rigid body docking of predicted models in Chimera.

Classes 1, 2, 4, and 7 showed two extra densities consistent with the size and shape of MlaC connected to the MlaD ring, and with quality sufficient to dock in AlphaFold models. In classes 3 and 5 only one density was clear. Class 6 shows only one density of a shape and size consistent with MlaC. Other densities observed were not of sufficient quality to interpret and may represent flexible MlaC molecules, or some misalignment in the maps. The maps from all classes, combined with our knowledge from high-resolution structures (PDB 6XBD, PDB 5UWA) suggest that MlaC can bind to any MlaD chain: A/A’ (in which the transmembrane helix is tightly associated with MlaE), B/B’ (in which the transmembrane helix is interacting with the C-terminus of the interfacial helix, IF1 of MlaE), or C/C’ (in which the transmembrane helix is interacting with the N-terminus of IF1).

## Data Availability

Cryo-EM map corresponding to class 1 has been deposited in the EMDB (EMD-29041) and raw micrographs in EMPIAR (XXXX). The AlphaFold2 models have been deposited in ModelArchive.org: MlaA-MlaC DOI: 10.5452/ma-5g2cp; MlaD-MlaC with 6:1 stoichiometry (DOI: 10.5452/ma-a56fx); MlaD-MlaC with 6:6 stoichiometry (DOI: 10.5452/ma-86bkv). Primary models for the MlaA-MlaC and MlaD-MlaC predictions can be found in the file upon Model Download and additional rank outputs can be found in the accompanying data file. Raw sequencing reads were deposited to the Sequence Read Archive under bioproject PRJNA849600 (accessions SRR19664419–29). Scripts to analyze reads resulting from deep mutational scanning can be found in the following GitHub repository: https://github.com/MaxabHaase/MlaC. Plasmids generated in this study have been deposited in Addgene (for accession numbers, see **Supplementary Table 3**).

## List of abbreviations

(OM): Outer membrane
(IM): Inner membrane
(PL): Phospholipid
(LPS): Lipopolysaccharide
(Mla): Maintenance of Lipid Asymmetry
(SDS): Sodium dodecyl sulfate
(EDTA): Ethylenediaminetetraacetic acid
(GFP): Green Fluorescent Protein

## Acknowledgments

We thank members of the Bhabha/Ekiert labs for helpful discussions. We thank James Chen, Juliana Ilmain, Fred Rubino, and Joe Sudar for critical reading and feedback on our manuscript. We thank Ian Henderson (University of Queensland) for providing anti-MlaD antibodies. We gratefully acknowledge the following funding sources: NIH R35GM128777 (DCE), PEW-00033055 (GB), pilot funding from the NYU Langone Health Antimicrobial-resistant Pathogen Program (DCE and GB), NIH T32 predoctoral training grant 5T32AI007180-39 (MRM). Screening and cryo-EM data collection were carried out at the NYU cryo-EM core facility. We thank Alice Paquette, Bill Rice, and Bing Wang for support in the NYU cryo-EM facility. The computational requirements for this work, including EM data processing, were supported in part by the NYU Langone High Performance Computing (HPC) Core’s resources and personnel. Illumina sequencing was performed by NYU Langone’s Genome Technology Center (RRID: SCR_017929). This shared resource is partially supported by the Cancer Center Support Grant P30CA016087 at the Laura and Isaac Perlmutter Cancer Center. Molecular graphics and analyses performed with UCSF Chimera, developed by the Resource for Biocomputing, Visualization, and Informatics at the University of California, San Francisco, with support from NIH P41-GM103311.

## Competing Interests Statement

The authors declare no competing interests.

## Supplementary File Legends

**Supplementary Data File 1. Deep mutational scanning top hit statistics**. The “top hits” for positions in MlaC that are most sensitive to mutation. A position in MlaC was considered a hit if 5 or more amino acid substitutions at that position resulted in a fitness decrease of more than one standard deviation below the average fitness of all mutations. Residues are grouped by their location on MlaC, including: SS (signal sequence), buried within the structure, in the lipid binding pocket, or exposed to the solvent. Included are the average fitness costs (between 2 replicates of deep mutational scanning) across all mutations for a residue and the standard deviation of this average. Additionally, point mutants assessed from this group to qualify the deep mutational scanning data are shown along with the average fitness cost for that specific mutation between the two replicates and the complementation spot plate result. Finally, MlaC residues that are predicted to interact with MlaD or MlaA based on our AlphaFold2 predictions (within 4 Å contact distance) are indicated.

**Supplementary Table 1: Cryo-EM data acquisition parameters**.

**Supplementary Table 2: Cryo-EM processing information. Supplementary Table 3: Plasmids used in this study**.

**Supplementary Table 4: Strain Table**

## Notes

### Competing Interest Statement

The authors have declared no competing interest.

