## Supplemental figures for "Protein-protein interactions in the Mla lipid transport system probed by computational structure prediction and deep mutational scanning"

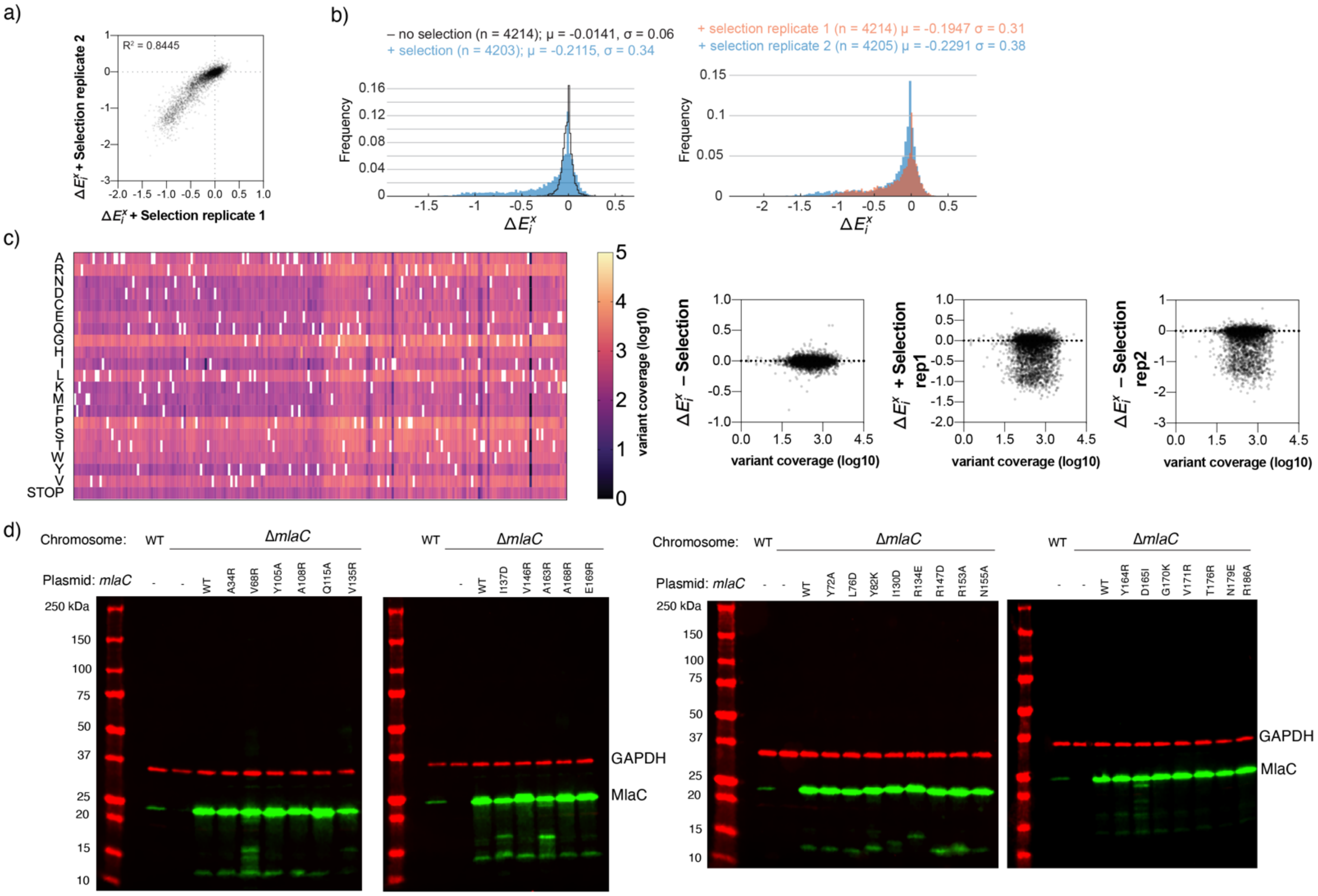

**Supplementary figure 1. Deep mutational scanning data statistics. a)** Relative fitness values of replicate 1 on X-axis and replicate 2 on Y-axis give a $R^2$ of 0.8445, validating the reproducibility of the experiment. **b)** Two histograms detailing the spread in cellular fitness impact (on x-axis) across all mutations. Y-axis represents frequency of occurrence; the relative number of mutations at a given cellular fitness impact can be observed. The left histogram details the average (between the two replicates) cellular fitness scores per mutation both with (blue) and without (black outline) selection. The average (μ) and standard deviation (σ) are shown for each. The right histogram shows the spread in cellular fitness impact for all mutations in the selection for replicate 1 (red) and replicate 2 (blue), again with the average (μ) and standard deviation (σ) shown for each. **c)** Heat map of coverage for each mutation made on MlaC. Overall, each mutant is at least represented 100 times. White rectangles represent the residues corresponding to the WT sequence of MlaC. Coverage vs fitness plots, indicate that apparent fitness is largely unaffected by variant coverage. **e)** Western blots of MlaC mutants used in genetic complementation experiments. Green bands correspond to MlaC (measured at 800 nm) and red bands correspond to the GAPDH loading control (measured at 680 nm).

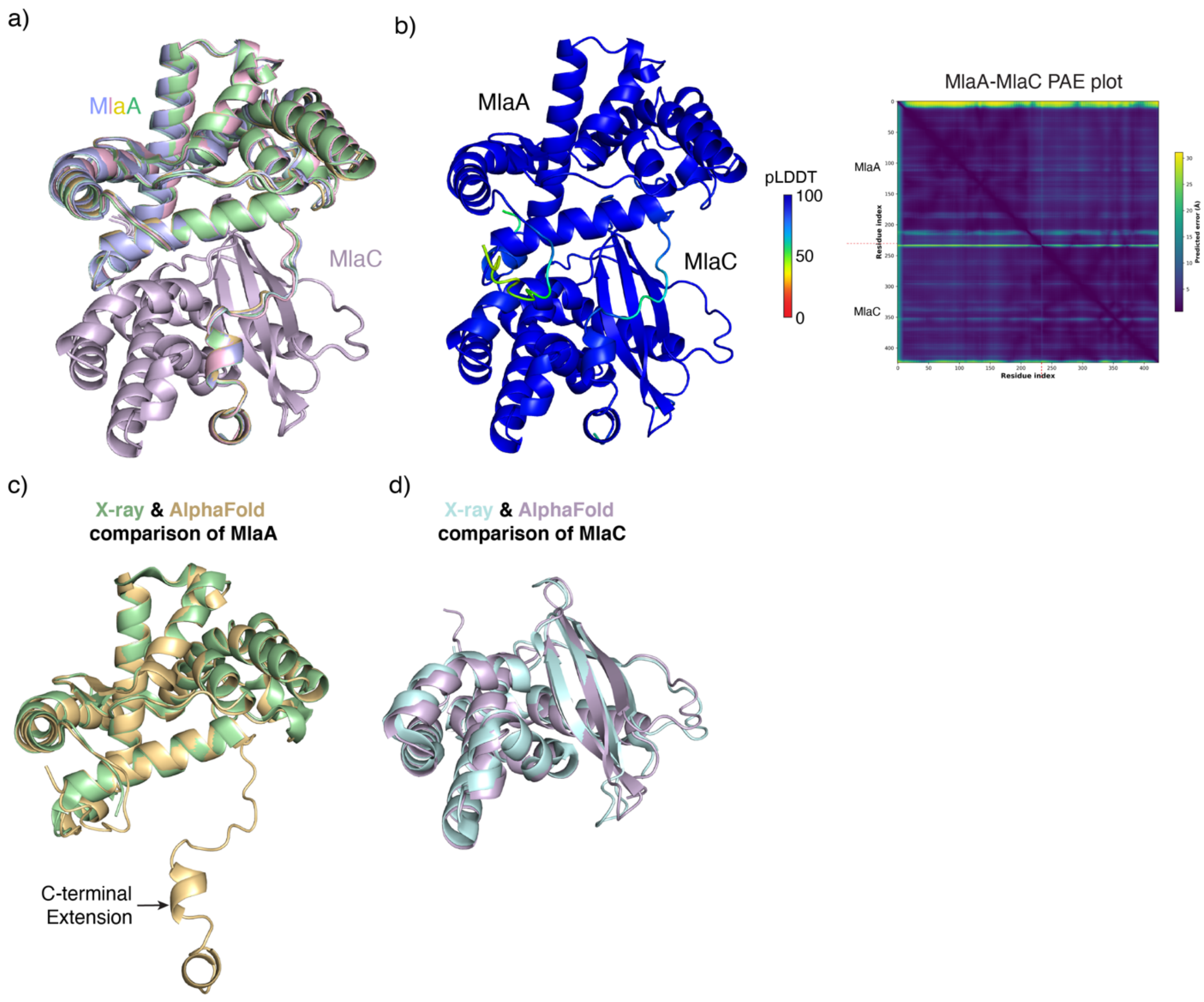


**Supplementary figure 2. Additional data for MlaC-MlaA predictions. a)** The 5 different 1:1 MlaC:MlaA AlphaFold Multimer predictions show a high similarity between each model (model 1, gold; model 2, green; model 3, blue; model 4, pink; model 5, cyan). Models have been aligned on MlaC, shown in purple. **b)** Confidence statistics for the MlaA-MlaC AlphaFold Multimer prediction are shown with pLDDT values colored as in the key on the predicted complex and the predicted alignment error (PAE) values between the two proteins plotted on a heatmap. **c)** Superimposition of MlaA from predicted model 1 (gold) and the crystal structure of MlaA (green; PDB 5NUO). **d)** Superimposition of MlaC from predicted model 1 (purple) and the crystal structure of MlaC (cyan; PDB code 5UWA).

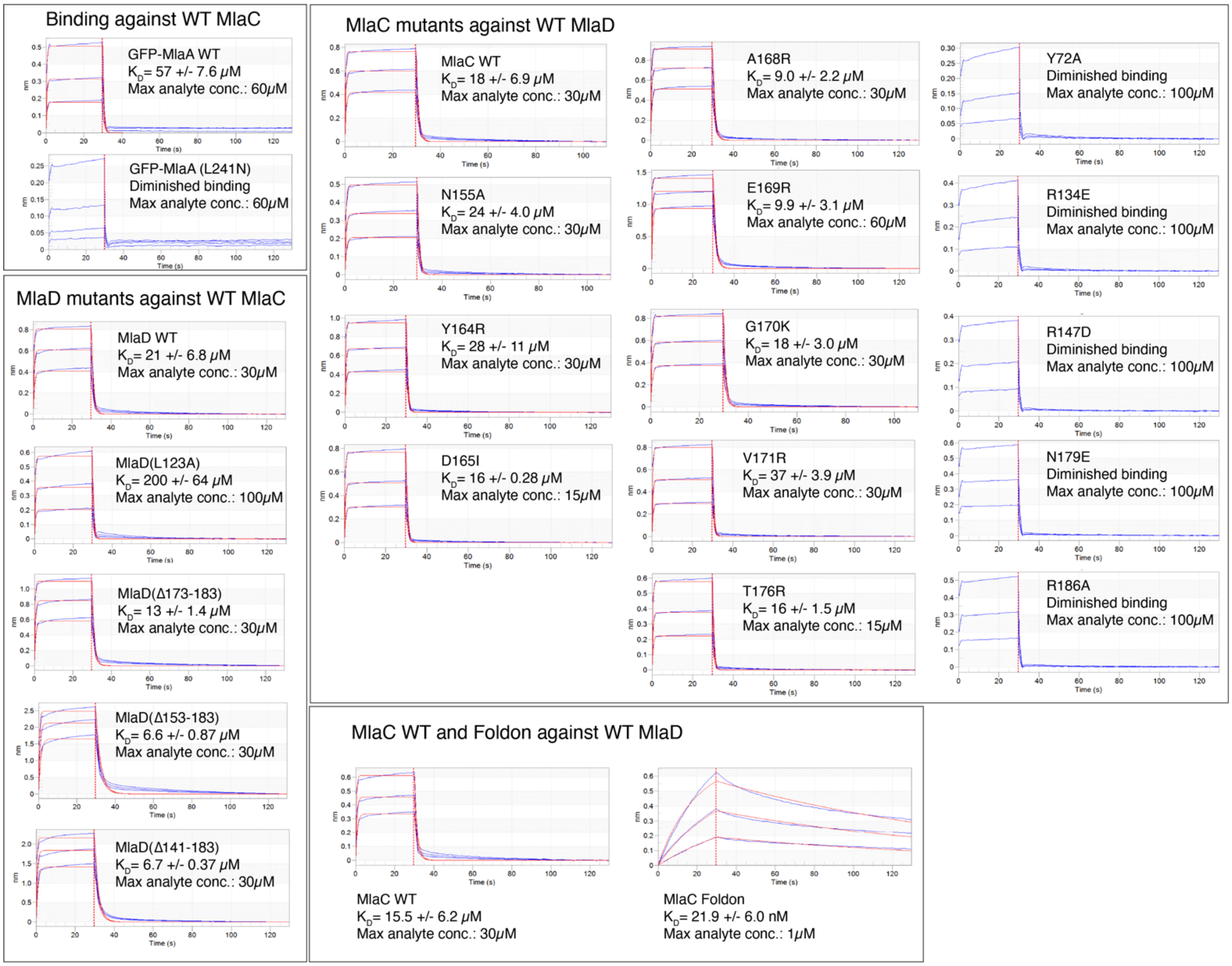


**Supplementary figure 3. Binding curve fits for biolayer interferometry experiments.** Curves for full titration (blue) and fits (red), for each binding interaction. A 1:1 binding model was used to generate a binding constant ($K_D$, μM) with an error between experiments (n=2).

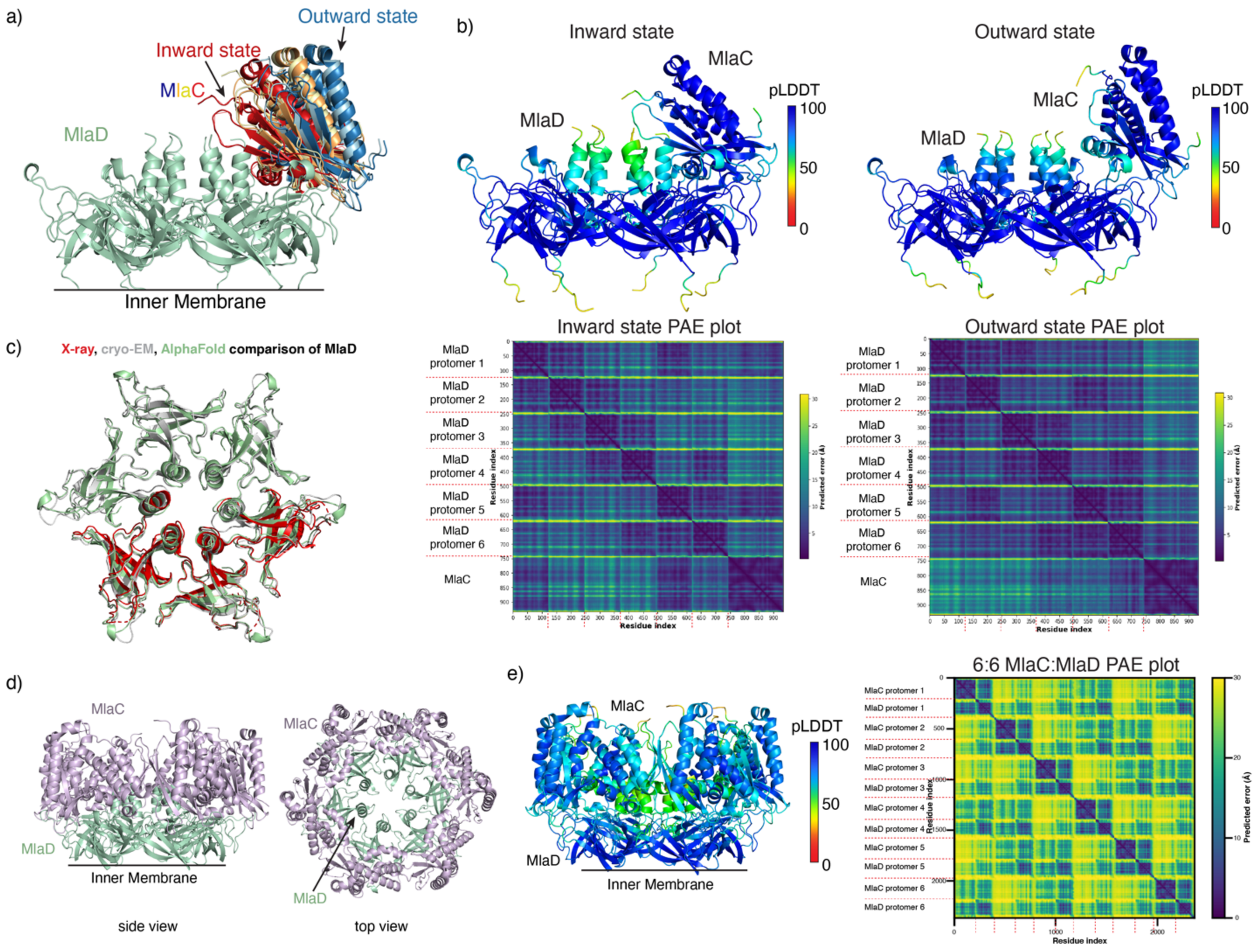


**Supplementary figure 4. Additional data for MlaC-MlaD predictions. a)** The 5 different 1:6 MlaC:MlaD predictions are shown with the 5 MlaC molecules colored from red to orange, yellow, light blue, and blue as they go from inward-facing to most outward facing states. Models are aligned on the MlaD ring. The inner membrane is indicated as a black line below MlaD. **b)** Confidence statistics for the inward and outward state AlphaFold2 predictions are shown. The pLDDT values are shown colored as in the key on the predicted models. The PAE values between chains in both predictions are plotted on a heatmap. **c)** Superposition of predicted MlaD from model 1 (green) with the crystal structure (red; PDB 5UW2), and the cryo-EM structure (gray; PDB 6XBD). **d)** The 6:6 MlaC:MlaD prediction; side and top facing views, with each MlaC colored in purple MlaD in green. One MlaC is observed to interact with the outer loop of one protomer and remain oriented roughly perpendicular to MlaD. The top five predicted models are consistent with each other. The inner membrane is indicated as a black line below MlaD. **e)** Confidence statistics for the MlaC-MlaD 6:6 AlphaFold2 prediction are shown with pLDDT values colored as in the key on the predicted complex and the predicted alignment error (PAE) values between the proteins plotted on a heatmap.

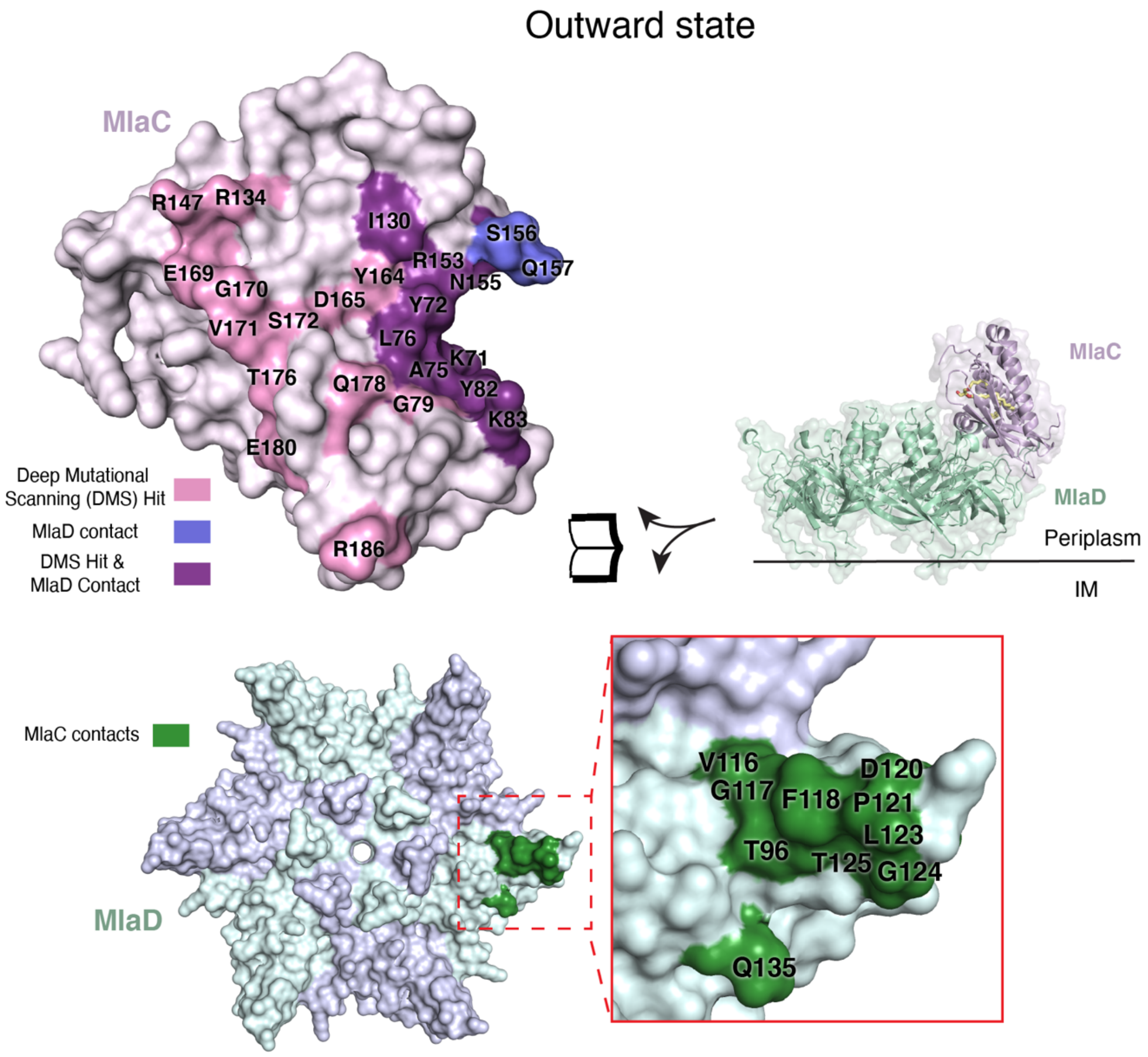


**Supplementary figure 5. Predicted protein-protein interactions of MlaC-MlaD in the outward state.** MlaC and MlaD are rotated as indicated and shown as molecular surfaces, highlighting interaction interfaces in the inward state. Residues in MlaC predicted to be within 4 Å distance of MlaD from AlphaFold2, shown to have reduced fitness from deep mutational scanning (DMS), or both, are mapped onto the structure as indicated in the key. Residues in MlaD predicted to be within 4 Å distance of MlaC from AlphaFold2 are mapped onto the structure as indicated in the key.

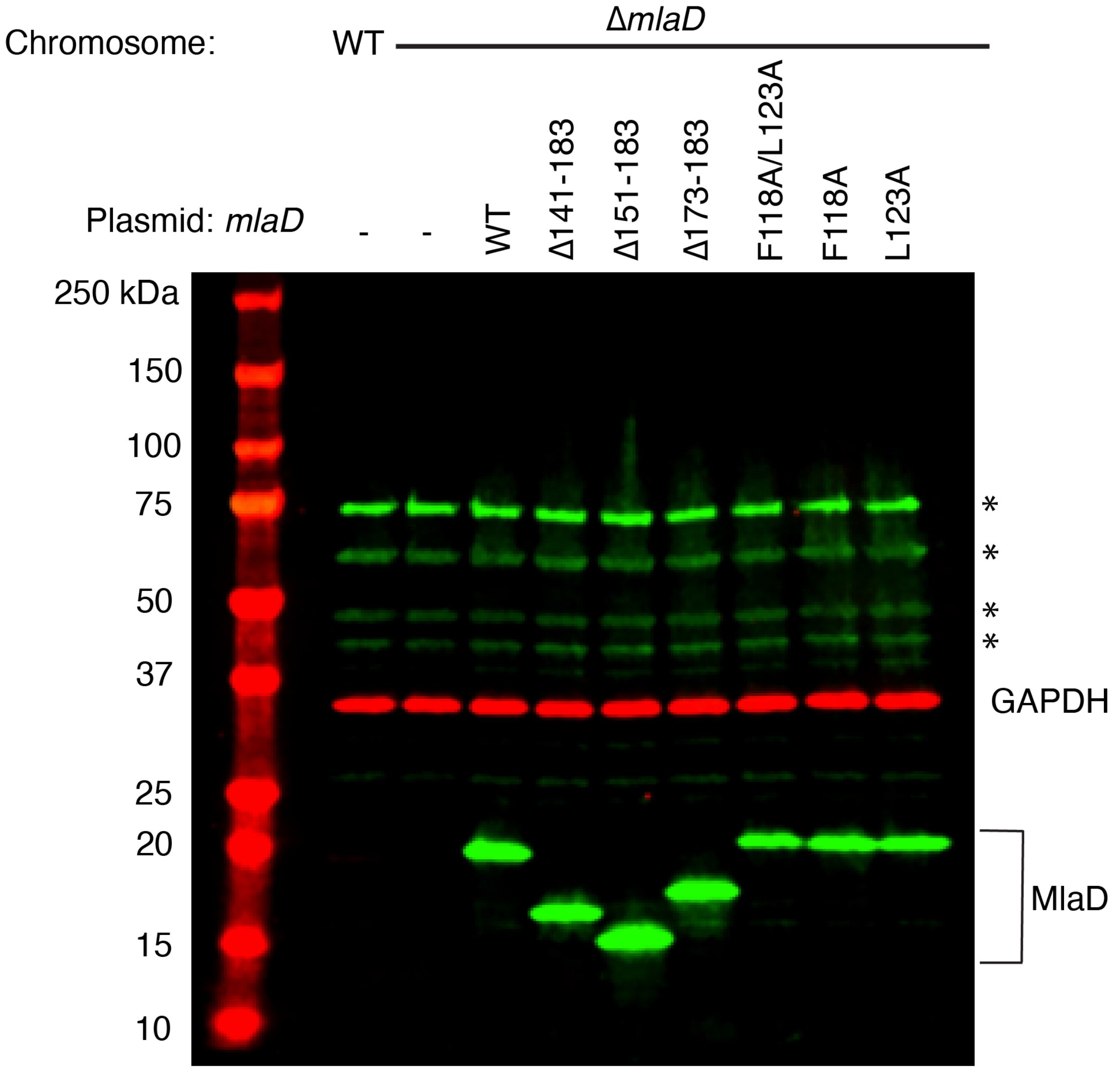


**Supplementary figure 6. Western blots for MlaD mutants. a)** Western blot again MlaD, corresponding to mutants used in genetic complementation experiments. Green bands correspond to MlaD (measured at 800 nm) and red bands correspond to the GAPDH as a loading control (measured at 680 nm). Asterisks correspond to non-specific bands present in all strains, including the Δ*mlaD* strain. Signal for MlaD in the WT strain is only marginally above background, and is highly over-expressed in complemented strains. These proteins are presumed to not be MlaD as the bands are also present in the *mlaD* KO strain sample.

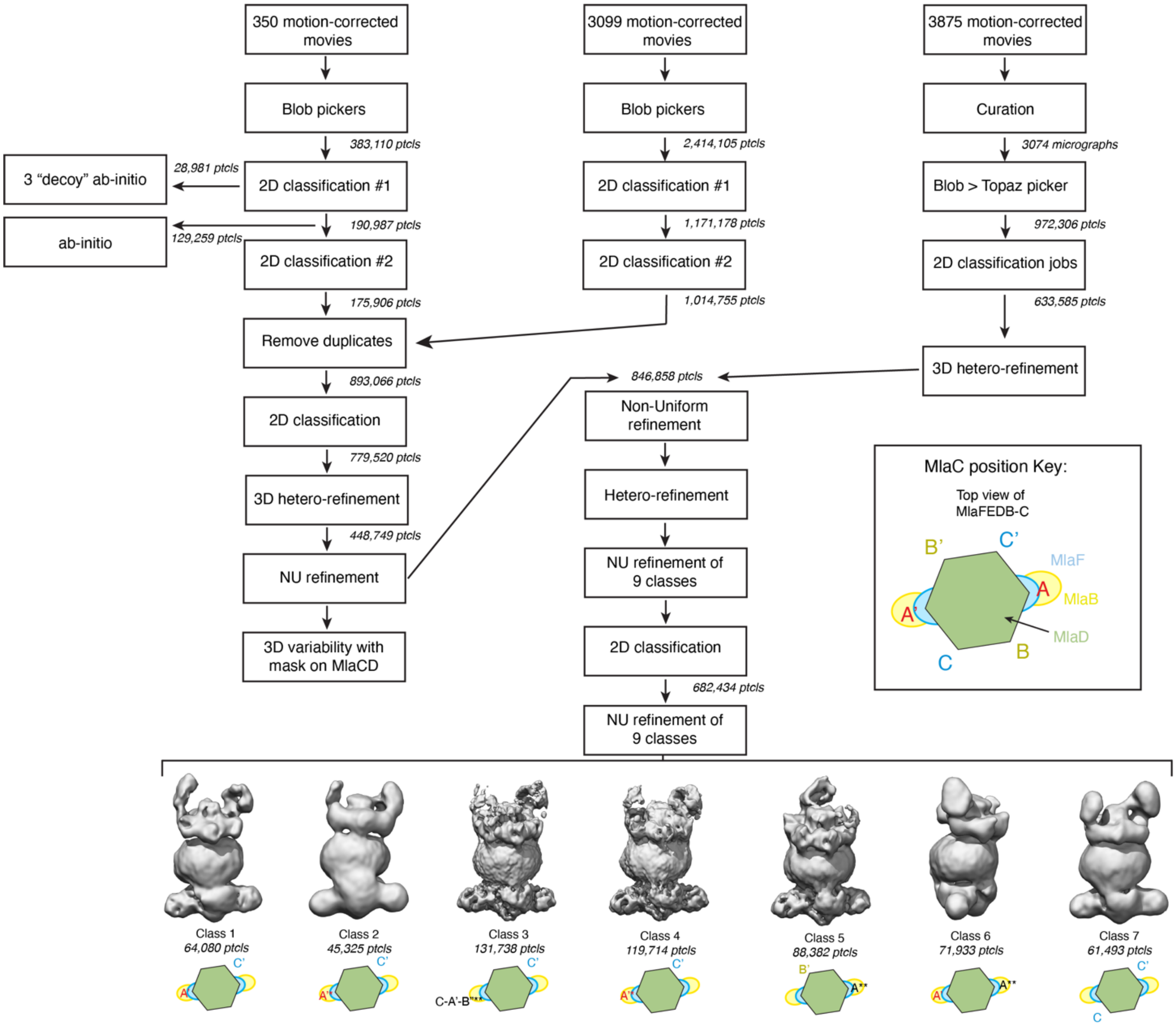


**Supplementary figure 7. Cryo-EM data processing workflow.** For the seven low-resolution classes obtained from heterogeneous refinement, a schematic shows which chain of MlaD is primarily interacting with MlaC. A schematic top view of MlaFEDB is represented with MlaD as a green hexagon, MlaF and MlaB in blue and yellow, respectively. * denotes weak density of a size and shape clearly consistent with MlaC. ** denotes messy density, which may either represent flexibility in MlaC binding, or misalignment in our maps, which we cannot decipher at this resolution. Based on a previous high-resolution structure (PDB 6XBD) (Coudray et al., 2020), Chains A and A’ correspond to the MlaD monomers in which the MlaD transmembrane helices are tightly interacting with MlaE and chains B and B’ are those observed to interact with the C-terminus of the interfacial helix IF1 helix of MlaE, and chains C and C’ interact closely with the N-terminus of this helix. Below each class is a simplified diagram showing the combinations of additional MlaC density observed in 3D classes generated.
