## Supplementary Table 1 & 2 for "Protein-protein interactions in the Mla lipid transport system probed by computational structure prediction and deep mutational scanning"

**Table 1 - Data acquisition**

| Microscope | Arctica |
| --- | --- |
| Voltage (kV) | 200 |
| Nominal magnification | 36,000 |
| Detector | K3 |
| Pixel size (Å/pix) | 0.548 (super-resolution) |
| No. of micrographs | 3,449 untilted and 3,875 tilted (40 deg) |
| No. of frames | 40 and 48 |
| Total dose (e-/Å^2^) | 56.70 and 50.78 |
| Exposure time (s) | 2.8 sec and 2.4 sec |
| Defocus range (µm) | 1.5-2.9 and 1.1-2.6 |

**Table 2- CryoEM processing information**

| Software | CryoSparc 3 |
| --- | --- |
| No. of selected particles | 846,858 |
| Total no. of particles in the final 7 classes | 582,665 |
| Box size (pixels) | 304 |
| Symmetry | C1 |
| Number of particles | 64,080 |
| Map sharpening B factor (Å^2^) | None |
| EMDB | XXXX |
| Coordinates | rigid-body docking of predicted models in Chimera |
