## Supplementary Table 3 for "Protein-protein interactions in the Mla lipid transport system probed by computational structure prediction and deep mutational scanning"

**Supplementary Table 3: Plasmids used in this study**

| **Plasmid ID** | **Features** | **Addgene ID** | **Source** |
| --- | --- | --- | --- |
| **General** | | | |
| pCP20 | FLP recombinase | *n/a* | *Cherepanov et al.*, 1995 |
| pBEL1194 | pBAD-derivative with stuffer | *n/a* | Invitrogen |
| **Complementation of MlaA mutants** | | | |
| pBEL2511 | MlaA(WT) | In progress | This study |
| pBEL2512 | MlaA(∆238-251) | In progress | This study |
| pBEL2513 | MlaA(∆227-251) | In progress | This study |
| pBEL2639 | MlaA(I241N) | In progress | This study |
| pBEL2640 | MlaA(L245N) | In progress | This study |
| pBEL2641 | MlaA(I248N) | In progress | This study |
| pBEL2644 | MlaA(D198N) | In progress | This study |
| pBEL2663 | MlaA(F223N) | In progress | This study |
| pBEL2664 | MlaA(L230N) | In progress | This study |
| pBEL2665 | MlaA(∆244-251) | In progress | This study |
| **Expression/purification of MlaA mutants** | | | |
| pBEL1092 | 6xHis-TEV-sfGFP | In progress | This study |
| pBEL2707 | 6xHis-TEV-sfGFP-MlaA (227-251) | In progress | This study |
| pBEL2753 | 6xHis-TEV-sfGFP-MlaA (227-251; I241N) | In progress | This study |
| pBEL2754 | 6xHis-TEV-sfGFP-MlaA (227-251; L245N) | In progress | This study |
| pBEL2755 | 6xHis-TEV-sfGFP-MlaA (227-251; I248N) | In progress | This study |
| pBEL2756 | 6xHis-TEV-sfGFP-MlaA(227-251; ∆238-251) | In progress | This study |
| pBEL2757 | 6xHis-TEV-sfGFP-MlaA(227-251; ∆244-251) | In progress | This study |
| **Complementation of MlaC mutants** | | | |
| pBEL2403 | MlaC(WT) | In progress | This study |
| pBEL2521 | MlaC(A34R) | In progress | This study |
| pBEL2522 | MlaC(A108R) | In progress | This study |
| pBEL2523 | MlaC(A163R) | In progress | This study |
| pBEL2524 | MlaC(A168R) | In progress | This study |
| pBEL2525 | MlaC(D165I) | In progress | This study |
| pBEL2526 | MlaC(E169R) | In progress | This study |
| pBEL2527 | MlaC(G170K) | In progress | This study |
| pBEL2528 | MlaC(I130D) | In progress | This study |
| pBEL2530 | MlaC(I137D) | In progress | This study |
| pBEL2531 | MlaC(L76D) | In progress | This study |
| pBEL2532 | MlaC(N155A) | In progress | This study |
| pBEL2533 | MlaC(N179E) | In progress | This study |
| pBEL2534 | MlaC(Q115A) | In progress | This study |
| pBEL2535 | MlaC(R134E) | In progress | This study |
| pBEL2536 | MlaC(R147D) | In progress | This study |
| pBEL2537 | MlaC(R153A) | In progress | This study |
| pBEL2538 | MlaC(R186A) | In progress | This study |
| pBEL2539 | MlaC(T116D) | In progress | This study |
| pBEL2540 | MlaC(T176R) | In progress | This study |
| pBEL2541 | MlaC(V68R) | In progress | This study |
| pBEL2542 | MlaC(V135R) | In progress | This study |
| pBEL2543 | MlaC(V146R) | In progress | This study |
| pBEL2544 | MlaC(V171R) | In progress | This study |
| pBEL2545 | MlaC(Y72A) | In progress | This study |
| pBEL2546 | MlaC(Y82K) | In progress | This study |
| pBEL2547 | MlaC(Y105A) | In progress | This study |
| pBEL2548 | MlaC(Y164R) | In progress | This study |
| **Expression/purification of MlaC mutants** | | | |
| pBEL1203 | 6xHis-TEV-MlaC | In progress | Ekiert *et al*, 2017 |
| pBEL2552 | 6xHis-TEV-MlaC(A168R) | In progress | This study |
| pBEL2553 | 6xHis-TEV-MlaC(D165I) | In progress | This study |
| pBEL2554 | 6xHis-TEV-MlaC(E169R) | In progress | This study |
| pBEL2555 | 6xHis-TEV-MlaC(G170K) | In progress | This study |
| pBEL2556 | 6xHis-TEV-MlaC(I130D) | In progress | This study |
| pBEL2559 | 6xHis-TEV-MlaC(L76D) | In progress | This study |
| pBEL2560 | 6xHis-TEV-MlaC(N155A) | In progress | This study |
| pBEL2561 | 6xHis-TEV-MlaC(N179E) | In progress | This study |
| pBEL2563 | 6xHis-TEV-MlaC(R134E) | In progress | This study |
| pBEL2564 | 6xHis-TEV-MlaC(R147D) | In progress | This study |
| pBEL2565 | 6xHis-TEV-MlaC(R153A) | In progress | This study |
| pBEL2566 | 6xHis-TEV-MlaC(R186A) | In progress | This study |
| pBEL2568 | 6xHis-TEV-MlaC(T176R) | In progress | This study |
| pBEL2572 | 6xHis-TEV-MlaC(V171R) | In progress | This study |
| pBEL2573 | 6xHis-TEV-MlaC(Y72A) | In progress | This study |
| pBEL2574 | 6xHis-TEV-MlaC(Y82K) | In progress | This study |
| pBEL2576 | 6xHis-TEV-MlaC(Y164R) | In progress | This study |
| **Expression/purification of MlaD mutants** | | | |
| pBEL1160 | 6xHis-TEV-MlaD | In progress | Ekiert *et al*, 2017 |
| pBEL1161 | 6xHis-TEV-MlaD∆141-183 | In progress | Ekiert *et al*, 2017 |
| pBEL1224 | 6xHis-TEV-MlaD∆153-183 | In progress | This study |
| pBEL1223 | 6xHis-TEV-MlaD∆173-183 | In progress | This study |
| pBEL1954 | 6xHis-TEV-MlaD(F118A/L123A) | In progress | This study |
| pBEL1909 | 6xHis-TEV-MlaD(F118A) | In progress | This study |
| pBEL1910 | 6xHis-TEV-MlaD(L123A) | In progress | This study |
| **Complementation of MlaD mutants** | | | |
| pBEL1195 | MlaF-MlaE-MlaD-MlaC-MlaB | In progress | Ekiert *et al*, 2017 |
| pBEL2766 | MlaF-MlaE-MlaD∆141-183-MlaC-MlaB | In progress | Ekiert *et al*, 2017 |
| pBEL1936 | MlaF-MlaE-MlaD∆153-183-MlaC-MlaB | In progress | This study |
| pBEL1935 | MlaF-MlaE-MlaD∆173-183-MlaC-MlaB | In progress | This study |
| pBEL1953 | MlaF-MlaE-MlaD(F118A/L123A)-MlaC-MlaB | In progress | This study |
| pBEL1933 | MlaF-MlaE-MlaD(F118A)-MlaC-MlaB | In progress | This study |
| pBEL1934 | MlaF-MlaE-MlaD(L123A)-MlaC-MlaB | In progress | This study |
| **Cryo-EM of MlaC-MlaFEDB** | | | |
| pBEL2319 | 6xHis-Foldon-(Gly-Ser-Linker)-MlaC | In progress | This study |
| pBEL1200 | MlaF-MlaE-(6xHis-MlaD)-MlaC-MlaB | 138526 | Ekiert *et al*, 2017 |
