## Supplementary Table 4 for "Protein-protein interactions in the Mla lipid transport system probed by computational structure prediction and deep mutational scanning"

**Supplementary Table 4: Bacterial strains used in this study**

| **Strain ID** | **Genotype** | **Source** |
| --- | --- | --- |
| BW25113 | “WT”; Δ(*araD-araB*)567 Δ(*rhaD-rhaB*)568 Δ*lacZ4787*(::*rrnB-3*) *hsdR514* *rph-1* | Coli Genetic Stock Center |
| JW3160 | BW25113 ∆*mlaD* (Keio background) | Baba 2006 |
| JW2343 | BW25113 *∆mlaA* (Keio background) | Baba 2006 |
| JW3159 | BW25113 *∆mlaC* (Keio background) | Baba 2006 |
| bBEL182 | BW25113 ∆*mlaD* | Coudray 2020 |
| bBEL465 | BW25113 *∆mlaA* | This study |
| bBEL464 | BW25113 *∆mlaC* | This study |
